# Notch controls the cell cycle to define leader versus follower identities during collective cell migration

**DOI:** 10.1101/2021.05.27.445572

**Authors:** Zain Alhashem, Dylan Feldner-Busztin, Christopher Revell, Macarena Alvarez-Garcillan Portillo, Joanna Richardson, Manuel Rocha, Anton Gauert, Tatianna Corbeaux, Victoria E Prince, Katie Bentley, Claudia Linker

## Abstract

Coordination of cell proliferation and migration is fundamental for life, and its dysregulation has catastrophic consequences, as cancer. How cell cycle progression affects migration, and vice-versa, remains largely unknown. We address these questions by combining *in silico* modelling and *in vivo* experimentation in the zebrafish Trunk Neural Crest (TNC). TNC migrate collectively, forming chains with a leader cell directing the movement of trailing followers. We show that the acquisition of migratory identity is autonomously controlled by Notch signalling in TNC. High Notch activity defines leaders, while low Notch determines followers. Moreover, cell cycle progression is required for TNC migration and is regulated by Notch. Cells with low Notch activity stay longer in G1 and become followers, while leaders with high Notch activity quickly undergo G1/S transition and remain in S-phase longer. We propose that migratory behaviours are defined through the interaction of Notch signalling and cell cycle progression.

## INTRODUCTION

The harmonious coupling of cell proliferation with migration is fundamental for the normal growth and homeostasis of multicellular organisms. A prominent consequence of its dysregulation is cancer. Primary tumours arise from uncontrolled cell proliferation, and the acquisition of migratory capacities leads to the formation of secondary tumours, the most common cause of cancer deaths. Metastatic cells can migrate collectively, which endows them with more aggressive behaviours (Nagai et al., 2020). Collective cell migration refers to the movement of a group of cells that maintain contact and read guidance cues cooperatively (Rørth, 2009). This mechanism has been studied in several contexts, such as wound healing, angiogenesis, and neural crest migration. However, how cell proliferation impacts collective cell migration, and vice-versa, remains largely unknown. The molecular signals that may couple these two fundamental processes remain equally unclear.

The NC is a mesenchymal cell population that arises early in development and migrates throughout the body giving rise to a variety of cell types (neurons, glia, pigment cells, etc.). Their stereotypical migratory behaviour (Gammill and Roffers-Agarwal, 2010) and similarity to metastatic cells (Maguire et al., 2015) makes the NC an ideal model to study the mechanisms of collective cell migration *in vivo*. Our previous work has shown that zebrafish trunk neural crest (TNC) migrate collectively forming single file chains (Richardson et al., 2016). One cell at the front of the chain, the leader, is the only cell capable of instructing directionality to the group, while follower cells trail the leader. This division of roles into leaders and followers has been observed in other collectively migrating systems (Theveneau and Linker, 2017). Moreover, histopathological studies from cancer samples and cell lines show clear morphological and molecular differences between the invasive front, leaders, and the lagging cells, followers (Pandya et al., 2017). One outstanding question from these studies is that of the signals which determine leader versus follower migratory identities.

Notch signalling is a cell-cell communication pathway that directly translates receptor activation at the membrane into gene expression changes. Notch receptors are activated by membrane-bound ligands of the Delta/Serrate/Lag2 family. Upon ligand binding, Notch receptors are cleaved by γ-secretases releasing its intracellular domain (NICD). Subsequently, NICD translocates to the nucleus, binds the CBF1/Su(H)/Lag-1 complex and initiates transcription (Bray, 2016). Among the direct Notch targets are members of the Hes gene family, which encode transcriptional repressors able to antagonize the expression of specific cell fate determinants and Notch ligands, generating a negative feedback loop in which cells with high Notch receptor activity downregulate the expression of Notch ligands, and cannot activate the pathway in their neighbours. Hence, adjacent cells interacting through the Notch pathway typically end up with either low or high levels of Notch activity and adopt distinct fates, a mechanism known as lateral inhibition (Lewis, 1998). Interestingly, Notch signalling has also been implicated in cell migration (Giniger, 1998; Leslie et al., 2007; Timmerman, 2004) and promotes invasiveness during cancer progression (Reichrath and Reichrath, 2012). Furthermore, lateral inhibition is implicated in the allocation of migratory identities during angiogenesis (Phng and Gerhardt, 2009), trachea formation in *Drosophila* (Caussinus et al., 2008) and in cell culture (Riahi et al., 2015). Whether Notch signalling plays a similar role in the context of mesenchymal cell migration is unknown. Notch signalling is required for NC induction (Cornell and Eisen, 2005) and its components and activity remain present in migrating NC (Liu et al., 2015; Rios et al., 2011). Nevertheless, the role of Notch during NC migration remains unclear (High et al., 2007; Mead and Yutzey, 2012; Vega-López et al., 2015).

The Notch pathway has not only been implicated in cell fate allocation, but it is also important for cell proliferation. Depending on the context, Notch can inhibit or promote cell cycle progression (Campos et al., 2002; Carlson et al., 2008; Devgan et al., 2005; Fang et al., 2017; Georgia et al., 2006; Mammucari et al., 2005; Nguyen et al., 2006; Nicoli et al., 2012; Noseda et al., 2004; Ohnuma et al., 1999; Park et al., 2005; Patel et al., 2016; Rangarajan et al., 2001; Riccio et al., 2008; Zalc et al., 2014). Indeed, Notch target genes include important cell cycle regulators as CyclinD1, p21 and MYC (Campa et al., 2008; Guo et al., 2009; Joshi et al., 2009; Palomero et al., 2006; Ronchini and Capobianco, 2001).

Using a combination of *in vivo* and *in silico* approaches we have herein established that differences in Notch activity between premigratory TNC select the leader cell. Cells with high levels of Notch signalling adopt a leader identity, while cells that lack Notch activity become followers. Our data show that a single progenitor cell in the premigratory area divides asymmetrically giving rise to a large prospective leader and smaller follower cell. We propose that this original small asymmetry generates differences in Notch activity between TNC that are thereafter enhanced by cell-cell communication through Notch lateral inhibition. Differences in Notch activity in turn drive distinct cell cycle progression patterns. Leader cells undergo the G1/S transition faster and remain in S-phase for longer than follower cells. Moreover, continuous progression through the cell cycle is required for TNC migration. Taken together, our results support a model in which the interaction between Notch and the cell cycle defines leader and follower migratory behaviours.

## RESULTS

### Notch signalling is required for TNC migration

NC cells are induced at the border of the neural plate early during development. Notch components are expressed in prospective NC at early stages, and Notch activity has been shown to be required for zebrafish NC induction (Cornell and Eisen, 2005). Our in-situ hybridization experiments show that Notch signalling components are retained by NC from induction throughout migration (Figure S1). To study the role of Notch signalling in NC development after induction, we first defined the developmental stage at which NC induction becomes independent of Notch signalling. To this end, we treated embryos with the γ-secretase inhibitor DAPT (Richter et al., 2017) and assessed NC induction by Crestin expression. Our results show that Notch inhibition impairs NC induction up to 11 hours post fertilization (hpf). Thereafter, NC establishment is independent of Notch signalling (Figure S2). Hence, all experiments were performed by altering Notch signalling at 12hpf, avoiding early defects in NC induction. Under Notch inhibition conditions, formation of all TNC derivatives is impaired (neurons, glia, and pigment; Figure 1A-F), suggesting that Notch activity is important in a process subsequent to induction yet prior to differentiation. Hence, we explored whether TNC migration is affected by Notch inhibition. Analysis of Crestin expression showed a reduction in both the number of TNC chains and their ventral advance in DAPT-treated embryos (Figure 1G-H), likely explaining the lack of TNC derivatives at later stages. We then asked whether these results are due to a delay or halt in migration. To this end, embryos were treated with DAPT for 6-12h and processed for Crestin expression. The number of migratory chains was decreased but as the embryos developed, new chains were formed, indicating that a blockade of Notch signalling delays TNC migration (Figure 1K). Comparable results were obtained by inhibiting Notch genetically in embryos where the dominant-negative form of Suppressor of Hairless is under the control of a heat shock element (hs:dnSu(H), Figure 1L; Latimer et al., 2005). We hypothesised that if Notch inhibition delays the onset of TNC migration, its overactivation might lead to an earlier onset of migration and an increased number of chains. To test this, we induced NICD expression by heat shock of hs:Gal4;UAS:NICD embryos (Scheer and Campos-Ortega, 1999). To our surprise, Notch gain and loss of function resulted in almost identical phenotypes, both showing similar reduction of TNC chain numbers (Figure 1L). Taken together, these results show that precisely regulated levels of Notch signalling are required for TNC migration.

**Figure 1.**
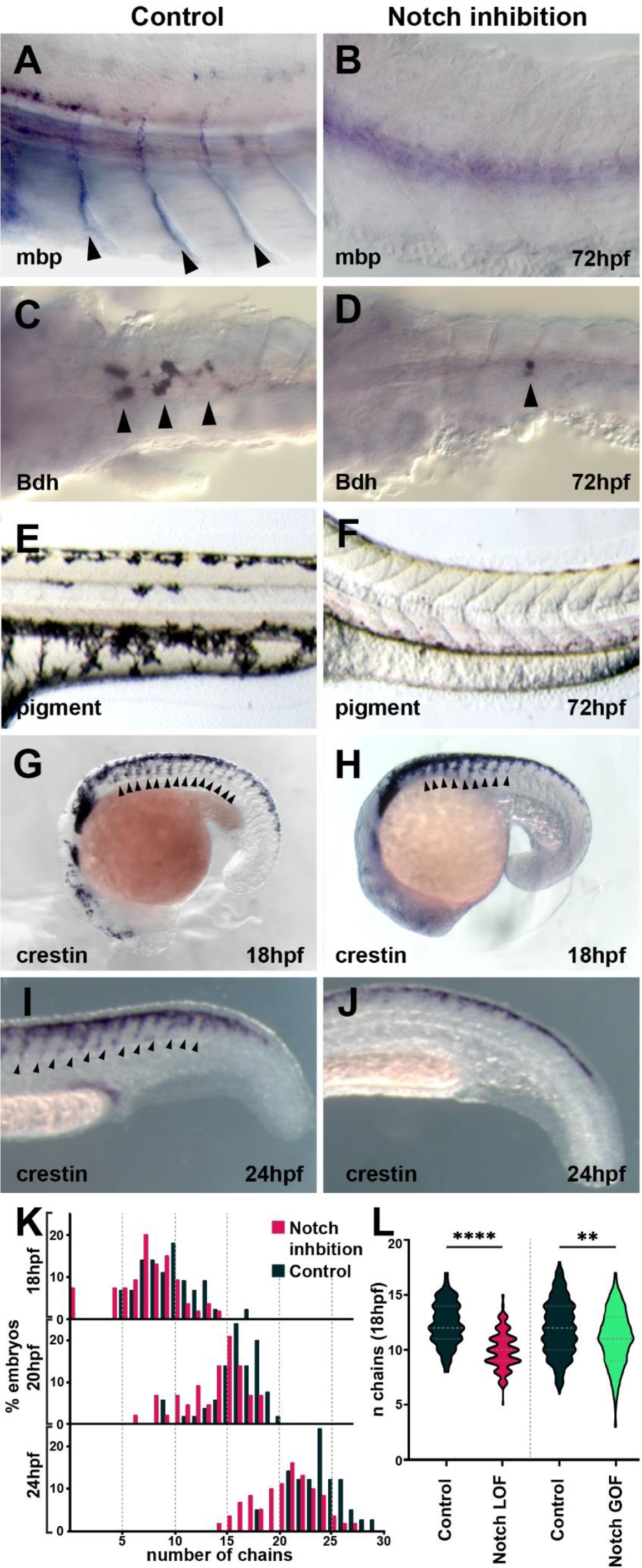
Notch signalling is required for TNC migration and derivatives formation. **A-B**. Glial marker mbp in situ hybridisation upon (A) control (DMSO; n=15) and (B) DAPT (n=20) treatment from 12hpf. **C-D**. Neuronal marker Bdh in situ hybridisation upon (C) control (DMSO; n=25) and (D) DAPT (n=18) treatment from 12hpf. **E-F**. Pigmentation upon (E) control (DMSO; n=40) and (F) DAPT (n=52) treatment from 12hpf. **G-H**. Crestin in situ hybridisation upon (G) control (DMSO) and (H) DAPT treatment from 12-18hpf. **I-J**. Crestin in situ hybridisation upon (I) control (DMSO) and (J) DAPT treatment from 12-24hpf. **K.** Quantification of migratory chain formation upon control (DMSO) and DAPT treatment from 12hpf to 18hpf (DMSO n=98; DAPT n=126), 20hpf (DMSO n=111; DAPT n=109) and 24hpf (DMSO n=42; DAPT n=61). **L.** Quantification of migratory chain formation in control (HS:Gal4; n=516), Notch LOF (HS:dnSu(H); n=220) and GOF conditions (HS:Gal4xUAS:NICD; n=142) heat shocked at 11hpf and analysed at 18hpf. Mann-Whitney U test, control vs LOF p<0.0001 ****, control vs GOF p=0.0020 **. Anterior to the left, dorsal top, except in C-D anterior left ventral view. Arrowheads indicate gene expression. All treatments were performed at 12hpf.

### *In vivo* Notch activity allocates TNC migratory identity

Interestingly, differences in Notch activity levels, established through lateral inhibition, have been shown to define distinct identities in model systems of collective migration. Hence, we hypothesised that Notch signalling may play a similar role allocating TNC migratory identity. To test this hypothesis, we performed live-imaging analysis (Figure 2–3 and Video S1-S2) and *in silico* modelling (Figure 3 and Video S3) of TNC migration under lack (inhibition and loss of function, LOF) or overactivation (gain of function, GOF) of Notch signalling. We first tested the effect of Notch inhibition by treating embryos with another γ-secretase inhibitor, Compound E (CompE; Richter et al., 2017). In these conditions, TNC remain motile with a single cell initiating the chain movement, but in contrast to control treatment (DMSO) the leader is unable to retain the front position, being overtaken by one or more followers (Figure 2A and C, and 3E; Video S1). The overtaking follower cell, in turn, is not always able to retain the front position and can be overtaken by cells further behind in the chain. This loss of group coherence corresponds with a reduction in ventral advance, with the majority of leader cells unable to move beyond the neural tube/notochord (NT/not; Figure 3D and F). Cells remain motile, accumulating at this location, some cells repolarise moving anterior or posteriorly, crossing somite boundaries and joining adjacent chains. Under these conditions, leader cells show a decreased speed and directionality, behaving as followers (Figure 3G-H). Similar results were observed when Notch inhibition was achieved genetically by driving overexpression of dnSu(H) through heat shock in the entire embryo (not shown; hs:dnSu(H) line). These data suggest that upon Notch inhibition all cells adopt a follower identity, establishing a homogeneous group that is unable to migrate. Notch signalling is important for the development of tissues surrounding TNC that act as substrate for migration, raising the possibility that Notch signalling does not act cell autonomously within TNC, with the observed phenotypes being a consequence of somite and/or neural tube malformations. However, this appears unlikely, as somite development (formation, patterning, and differentiation) and neuron formation are not affected by Notch inhibitors at the axial level analysed (Figure S3). Next, we directly tested whether Notch signalling is autonomously required in TNC, by inhibiting Notch activity exclusively in the NC at the time of migration. To this end, a new UAS:dnSu(H) line was generated and crossed to Sox10:Kalt4 fish (Alhashem et al., 2021). In these embryos all NC express Gal4 fused to the oestrogen receptor binding region (Gal4-ER) and are fluorescently labelled by nuclear-RFP. Under normal conditions, Gal4-ER is maintained inactive in the cytoplasm whilst upon addition of tamoxifen, Gal4-ER is translocated to the nucleus activating transcription from the UAS:dnSu(H) transgene (Figure S4). We found that autonomous inhibition of Notch signalling in NC phenocopies the chemical inhibition. Leader cells are unable to retain the front position, being overtaken by followers, and ventral advance is reduced with cells accumulating at the NT/not boundary (Figure 2D and 3D-F; Video S2). Moreover, leader cells adopt followers’ migratory parameters, showing decreased speed and directionality (Figure 3G-H). Together these data show that Notch activity is autonomously required in TNC for identity allocation. In the absence of Notch signalling, a homogenous group of followers is established.

**Figure 2.**
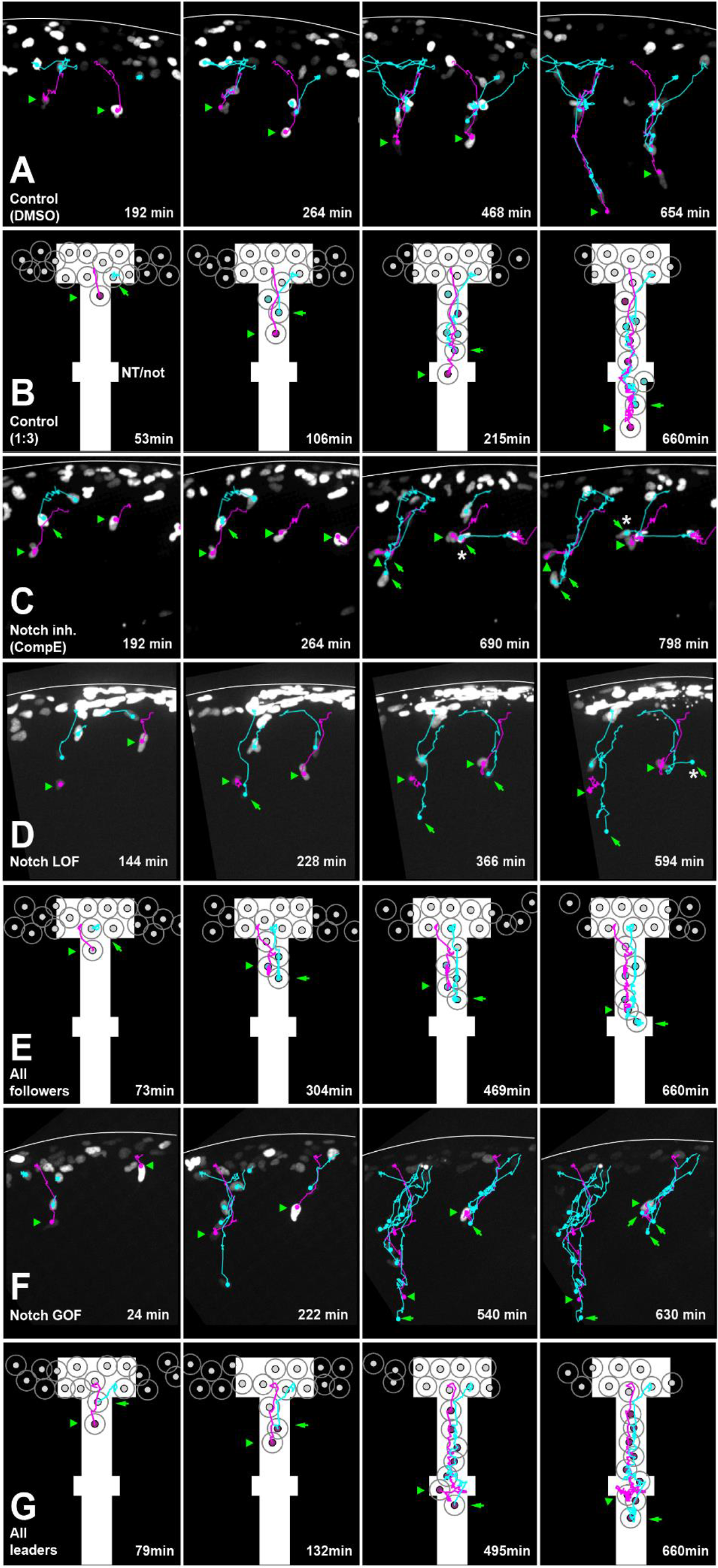
Notch activity allocates TNC migratory identity. **A.** Selected frames from in vivo imaging of Sox10:Kalt4 control (DMSO treated) embryos. **B.** Selected frames from control simulation with 1:3 leader/follower ratio. **C.** Selected frames from in vivo imaging under Notch inhibited condition, Sox10:Kalt4 embryos treated with CompE. **D.** Selected frames from in vivo imaging of Notch LOF condition, Sox10:Kalt4; UAS:dnSu(H) embryos. **E.** Selected frames from all followers simulation. F. Selected frames from in vivo imaging of Notch GOF condition Sox10:Kalt4; UAS:NICD embryos. **G.** Selected frames from all leaders simulation. Magenta tracks and green arrowheads indicate leaders; green arrows and cyan tracks followers. Asterisks indicate cells crossing somite borders. White lines in A, C, D and F mark the dorsal midline of the embryo. NT/not: neural tube/notochord boundary. Anterior to the left, dorsal up. Time in minutes.

**Figure 3.**
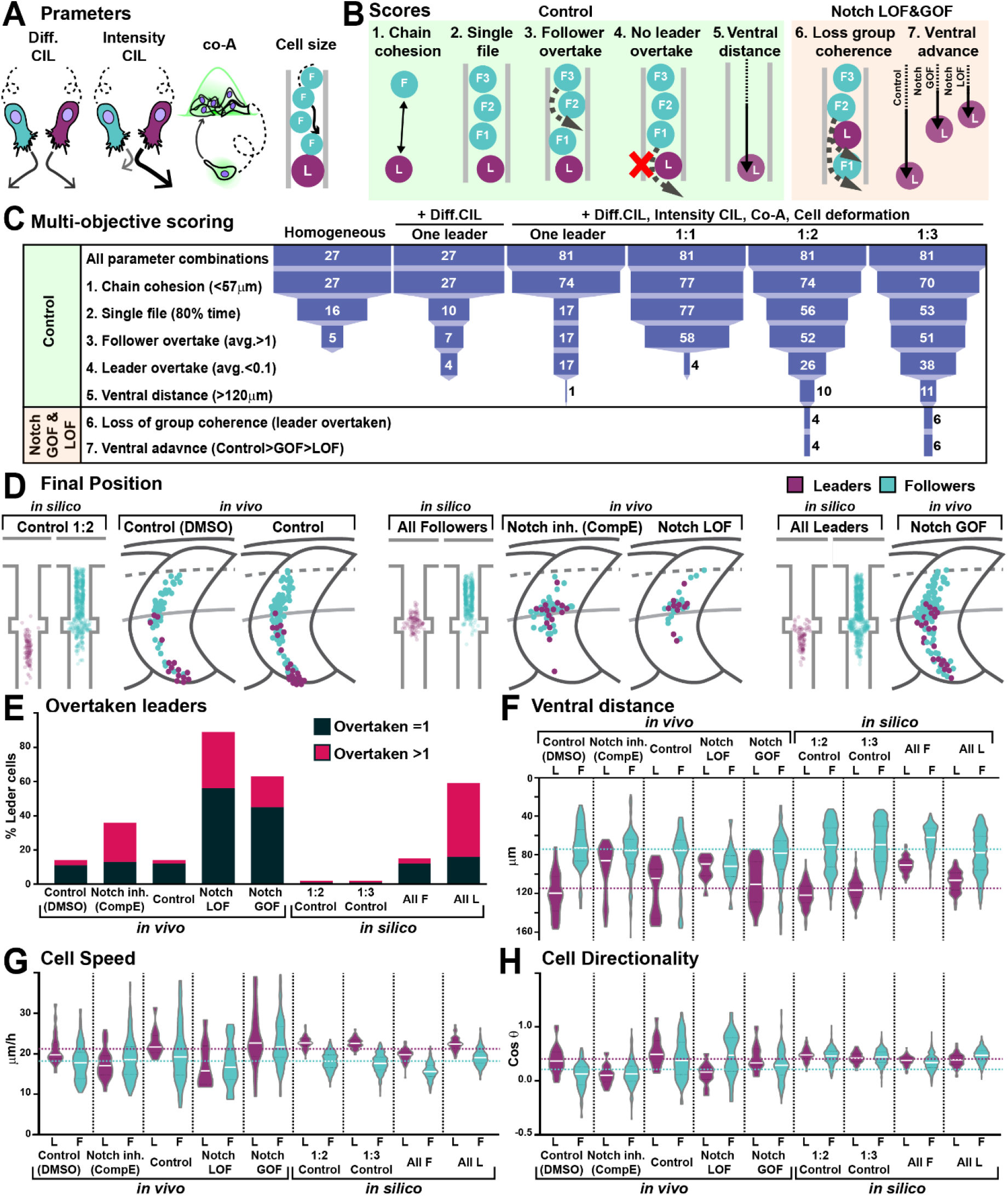
*In-silico* modelling predicts that more than one leader is required for TNC migration. **A.** Schematics of model parameters. **B.** Schematics of simulation scores. **C.** Depiction of parameter space analysis showing the number of parameter sets that fulfilled each score when different variables were tested. One leader, chains with a single leader cell. 1:1, 1:2 and 1:3 refer to leader/follower ratios. **D.** Final position of each cell in model simulations and in vivo experiments under different conditions. In silico results depicted in confined pathway, in vivo data graphed in model embryo, somites contour and dorsal midline dark grey lines, edge of the premigratory area dashed lines, and NT/not boundary light grey lines. Anterior left, dorsal up. **E.** Quantification of leader overtaking events in vivo and in silico. **F.** Quantification of the ventral advance of cells in vivo and in silico. **G.** Quantification of cell speed in vivo and in silico. **H.** Quantification of cell directionality in vivo and in silico. Leader cells in magenta, followers in cyan. Magenta dotted line mark average of control leaders, cyan line of control followers in F, G and H. Full statistical analysis in Table S1.

In view of these results, we hypothesised that a homogeneous group of leaders might form upon Notch overactivation. Using a similar strategy, Notch overactivation was induced in the whole embryo (not shown, hs:Gal4;UAS:NICD; Scheer and Campos-Ortega, 1999), or exclusively in NC (Sox10:Kalt4;UAS:NICD) and migration was analysed by live-imaging. Similar results were obtained in both experimental conditions: group coherence is lost, leader cells are overtaken by followers, and ventral advance is impaired (Figure 2F and 3D-F; Video S2). Interestingly, in Notch GOF conditions follower cells adopt leaders’ characteristics, moving with increased speed, but all cells in the chain follow less directional trajectories, which hinders the ventral advance of the group (Figure 3G-H). Our *in vivo* analysis show that Notch signalling is autonomously required in TNC for identity allocation. TNC with high Notch levels become leaders while cells with low Notch activity migrate as followers, suggesting that Notch-mediated lateral inhibition is a possible mechanism for identity allocation.

### *In-silico* modelling predicts that more than one leader is required for TNC migration and cell size to be a key difference between leader and follower cells

Our *in vivo* experiments show that both Notch inhibition and overactivation hinder TNC migration due to the establishment of homogeneous populations that lack group coherence. To gain a better understanding of these paradoxical results we developed a discrete element model of TNC migration. Cells were simulated as 2D particles moving into a constrained space and endowed with intrinsic motility. Four variables control cell movement in the model: contact inhibition of locomotion (CIL) and co-attraction (co-A) define movement directionality and group cohesion, while volume exclusion regulates cell overlap, intuitively understood as cell size (Figure 3A). Finally, an element of noise (zeta) is added to the cell’s trajectory. A multi-objective scoring system, based on *in vivo* measurements, was developed to evaluate how close simulations with different underlying mechanisms matched chain behaviours. The scores were: 1. chain cohesion, a maximum distance of 57μm is allowed between adjacent cells, 2. single file migration for at least 80% of the simulation 3. followers undergo rearrangements, while 4. leaders retain the front position, and 5. the chain should advance to the end of the migratory path (Figure 3B). Using this analysis and a parsimonious modelling approach, we attempted to match *in vivo* TNC migration with the simplest form of the model and only added complexity incrementally, in an effort to find the minimal set of predicted mechanisms required. We first simulated chains composed of homogeneous cells and systematically covaried all parameters. We found no parameter combination able to match all scores, confirming our previous findings that cell heterogeneity is required for TNC migration (Figure 3C; Richardson et al., 2016). Evidence from other systems (Astin et al., 2010; Bentley et al., 2014; Parkinson and Edwards, 1978; Theveneau and Mayor, 2013) led us to hypothesise that differences in the CIL response between cells may be at play. Thus, we simulated chains in which only cells of different identities present CIL (Diff CIL). These simulations match several scores, but chains are unable to reach the end of the migratory path (Figure 3C and Video S3). Next, we varied CIL intensity, co-A and cell size (volume exclusion) for leader cells. Interestingly, the model is only able to recapitulate control conditions when the difference between leaders and follower is maximal for all variables. Nevertheless, it is unable to recapitulate Notch GOF and LOF phenotypes (Figure 3C).

Our *in vivo* data show that differences in Notch signalling establishes migratory identities and we hypothesised that lateral inhibition could be the mechanism at play. To explore how different outcomes of lateral inhibition may impact TNC migration, different ratios of leader/follower cells were simulated. We first tested a 1:1 ratio, surprisingly this chain architecture over-migrates, moving beyond the end of the pathway (Figure 3C; Video S3). Interestingly, we found that several parameter combinations from the 1:2 and 1:3 leader/follower ratios were able to recapitulate *in vivo* control condition, as well as the loss of group coherence and ventral advance observed in Notch GOF (all leader simulation) and LOF (all follower simulation; Figure 2B, E and G and 3C-H; Video S3). In these simulations, the six parameter combinations that match all *in vivo* scores had followers at the low setting, while leaders’ CIL intensity took medium or high values, cell size medium or low values and co-attraction took on all three levels; but all these parameter combinations endow leader cells with enhanced migratory behaviour. Taken together, our *in-silico* data confirms our previous conclusion that TNC chains are a heterogeneous group. Remarkably, it also predicts that TNC chains are formed of leaders and followers in a 1:2 or 1:3 ratio, in which leaders are larger and better migratory cells than followers.

### Leader cells arise from the asymmetric division of a progenitor cell

Cell size is a prominent characteristic distinguishing leader from follower cells. Leaders are almost twice as big as followers during migration and this difference is evident before migration initiation (Richardson et al., 2016), suggesting that size disparity arises at birth or shortly thereafter. Interestingly, differential cell size emerged as an important parameter in our in silico analysis, contributing to more realistic leader/follower coordination behaviours. To understand the origin of these size differences we investigated whether leader and follower cells share a common progenitor, and at which point differences in size become apparent. To this end we imaged FoxD3:mCherry;H2aFVA:H2a-GFP embryos. The FoxD3:mCherry reporter (Lukoseviciute et al., 2018) labels NC from early stages and allows us to define TNC identity at later stages by their migratory position. Moreover, the nuclear marker H2aFVA:H2a-GFP (Pauls et al., 2001) was used to track single cells and their divisions. Tracking analysis shows that the asymmetric division of a single progenitor cell in each body segment gives rise to a larger cell that becomes a leader (102 ± 20 μm^2^), and a smaller sibling that migrates as follower (72 ± 9 μm^2^; Figure 4A and C). In contrast, all other progenitors divide symmetrically giving rise to two followers (87 ± 27 μm^2^; Figure 4B and D). We also noticed that the leader progenitors’ divisions are spatially restricted to the anterior quarter of the premigratory area in each segment, while the followers’ progenitor divisions take place across the premigratory area (Figure 4E).

**Figure 4.**
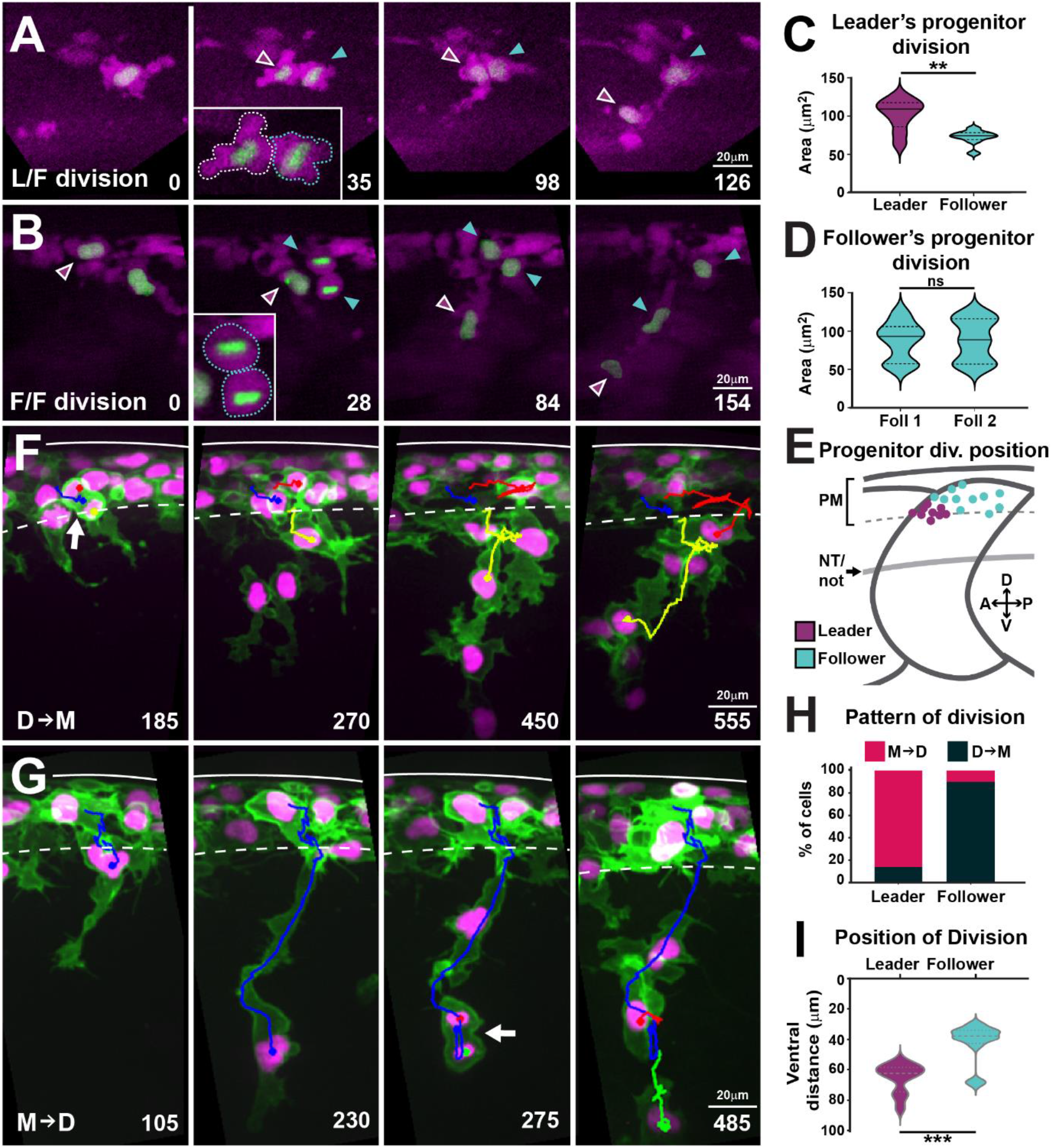
Leaders arise from the asymmetric division of a progenitor cell and present characteristic division patterns. **A.** Selected frames from in vivo imaging of leaders’ progenitor division in FoxD3:mCherry; H2aFVA:H2a-GFP embryos at the level of somite 7. **B.** Selected frames from in vivo imaging of followers’ progenitor division in FoxD3:mCherry; H2aFVA:H2a-GFP embryos at the level of somite 7. **C.** Area of leaders’ progenitor daughter cells (n=9 cells, 7 embryos; Mann-Whitney U test, p=0.0056). **D.** Area of followers’ progenitor daughter cells (n=10, 4 embryos; Mann-Whitney U test, p>0.9999). **E.** Position of progenitors’ divisions on model embryo (leaders n=9, 7 embryos; followers n=10, 4 embryos). PM: premigratory area; NT/not: neural tube/notochord boundary. **F.** Selected frames showing the D→M division pattern from 16-28hpf in vivo imaging of a Sox10:mG embryo. Blue before-, yellow and red after-division. Arrow indicates division position. **G.** Selected frames showing the M→D division pattern from 16-28hpf in vivo imaging ofa Sox10:mG embryo. Labelling as in F. **H.** Quantification of leaders’ (n=21, 7 embryos) and follower’s division patterns (n=43, 7 embryos). Red: M→D, black: D→M. **I.** Quantification of division positions (n as in C; Mann-Whitney U test, p=0.0002). Time in minutes. Leaders in magenta, followers in cyan. Anterior left, dorsal top.

We then reasoned that leader cells, being bigger, may undergo the next division in a shorter time span than follower cells and in consequence, mitotic figures would be observed at different, but consistent, positions in their trajectory. Indeed, we found two different patterns of divisions in respect to migration: i) cells that first <u>D</u>ivide and then <u>M</u>igrate (D→M), or ii) cells that first <u>M</u>igrate and then <u>D</u>ivide (M→D; Figure 4F-G; Video S4). Interestingly, we found that the patterns of cell division correlate with cell identity. The majority of leader cells divide during migration (M→D: 86%), while the bulk of follower cells divide before migration initiation (D→M: 90%, Figure 4H). These patterns result in leader and follower cells dividing at distinct positions, 74% of leaders divide at the NT/not boundary (65.3 ± 9.6 μm), while 85% of followers divide mostly within the premigratory area or in the dorsal-most region of the somite (42 ± 12.4 μm; Figure 4I). Together, these results show that leader cells arise from the asymmetric division of a progenitor. Thereafter, leader and follower cells show distinct locations and patterns of division, suggesting that leader and follower cells progress asynchronously through the cell cycle, which may influence their migratory behaviour.

### Cell cycle progression is required for TNC migration

To test the role of cell cycle progression in TNC migration directly, we used inhibitory drugs. The S-phase inhibitor Aphidicolin blocks over 94.7 ± 4.5% of mitotic figures after 3h of treatment, while the G2/M inhibitor Genistein prevents 90 ± 10% of divisions within 6h, but neither treatments affect NC induction (Figure S5). Inhibition of cell cycle progression by either of the treatments resulted in reduced numbers of migratory chains and decreased ventral advance (Control 19 ± 2, Genistein 10 ± 3, Aphidicolin 6 ± 2 chains; Figure 5A-H). This result was not due to the loss of cell motility, as premigratory TNC cells actively extend protrusions and move along the antero-posterior axis but are unable to migrate ventrally (Video S5). Importantly, these effects were not a consequence of cell death or the permanent impairment of motility, as TNC re-initiate migration and form new chains upon drug withdrawal (Figure 5G and H); showing that active cell cycle progression is required for migration. Next, we directly analysed TNC cell cycle progression *in vivo*. To this end, we imaged Sox10:FUCCI embryos (Rajan et al., 2018), in which TNC nuclei are RFP-labelled during G_1_ and GFP-labelled during S and G_2_. Tracking analysis show differential cell cycle progression, with most leader cells initiating migration in S-phase (79%), while followers start movement during G_1_ (77%; Figure 5I and J; Video S6). These results show that cell cycle progression is required for migration and that leader and follower cells initiate movement at different points of the cell cycle, suggesting an intimate connection between cell growth and movement.

**Figure 5:**
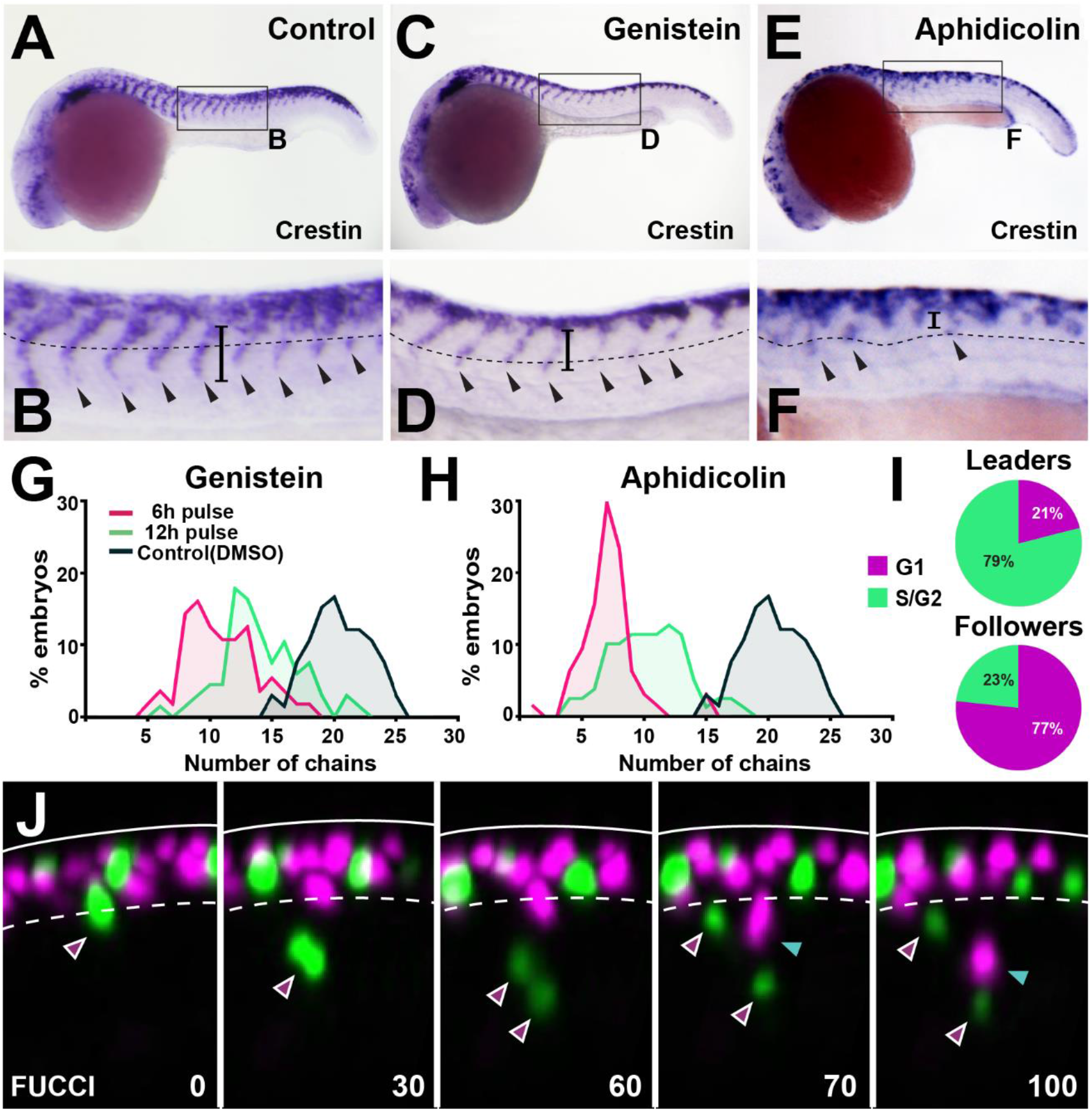
Cell cycle progression is required for TNC migration. **A, C and E**. Crestin in situ hybridisation upon (A) DMSO, (C) Genistein or (E) Aphidicolin treatment from 12-24hpf. **B, D and F**. Enlargement of areas indicated by boxes in (A, C, E). Dotted line marks NT/not boundary, arrowheads migratory chains and vertical line the chain length. **G-H**. Frequency distribution of migratory chains upon control (DMSO; n=66), (G) Genistein (12h pulse, n=56; 6h pulse, n=67) or **(H)** Aphidicolin (12h, n=64; 3h, n=79). **I.** Cell cycle phase at migration initiation for leaders (n=38, 4 embryos) and followers (n=43, 4 embryos). **J.** Selected framed from in vivo imaging of Sox10:FUCCI. Time in minutes. Solid line marks dorsal midline, dotted line the premigratory area. Magenta arrowheads indicate leader and its daughters. Green arrowheads indicate followers.

### Leader and follower cells progress through the cell cycle at different rates

Next, we studied TNC cell cycle progression in detail. First, we asked whether leaders and followers differ in the total length of their cell cycle. Measurements of the time span between two consecutive mitoses showed no significant differences in the total length of the cell cycle between leaders and followers (13.6 ± 1.2 and 13.3 ± 1.4 h respectively; Figure 6B). Next, we examined the length of each phase of the cell cycle by imaging the characteristic nuclear labelling pattern of the PCNA-GFP fusion protein (Leung et al., 2012). Sox10:Kalt4 embryos, in which all NC can be recognized by nuclear RFP expression, were injected with PCNA-GFP mRNA and live imaging was performed. PCNA-GFP shows uniform nuclear GFP labelling during G_1_, intense fluorescent nuclear puncta characterise the S-phase, these puncta dissipate during G_2_ restoring homogeneous nuclear fluorescence, at the onset of mitosis PCNA is degraded and TNC are recognized solely by nuclear RFP (Figure 6A; Video S7). In these embryos, leader cells initiate migration during S-phase and followers in G_1_, confirming our FUCCI results and establishing that PCNA overexpression does not introduce artefacts to cell cycle progression (Figure S6). Using this tool, we measured the length of the cell cycle phases in TNC.

**Figure 6.**
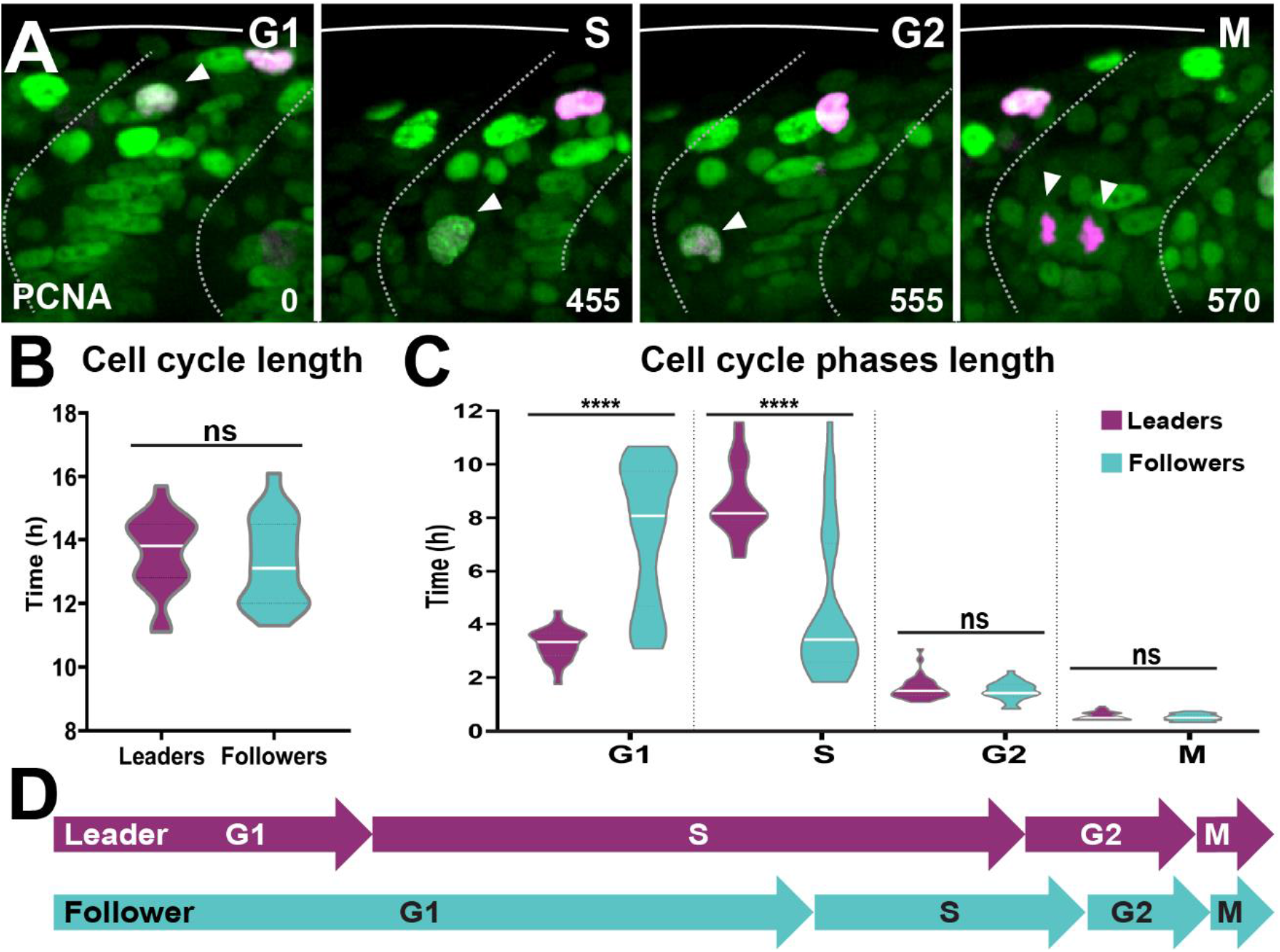
Leader and follower cells progress through the cell cycle at different rates. **A.** Selected frames from in vivo imaging from 16-28hpf of Sox10:Kalt4 embryos injected with PCNA-GFP mRNA. White arrow points to cycling. Time in minutes. **B.** Quantification of the cell cycle total duration in leaders (n=20, 7 embryos) and followers (n=19, 7 embryos; Unpaired t test, p=0.5240). **C.** Quantification of the cell cycle phases duration in leaders (G_1_ n=45, S n=44, G2 n=33 and M n=32, 11 embryos) and followers (G_1_ n=50, S n=48, G2 n=33 and M n=34, 11 embryos). Brown-Forsythe and Welch ANOVA tests, G_1_ p<0.0001, S p<0.0001, G2 p=0.9997, M p=0.9231. **D.** Schematic representation of the cell cycle phases durations.

We found striking differences in the time spent in G_1_- and S-phase between leader and follower cells. Leaders present a short G_1_ (3.2 ± 0.6h) but remain for twice as long in S-phase (8.7 ± 1.3h). Followers, on the other hand present the opposite distribution, remaining for twice as long in G_1_ (7.4 ± 2.7h) than in S-phase (4.6 ± 2.8h; Figure 6C-D). No significant differences were observed in the length of G_2_ (leaders 1.6 ± 0.4 h; followers 1.5 ± 0.3h) or M (leaders 0.6 ± 0.1h; followers 0.5 ± 0.1h). These data show that leader and follower cells present marked differences in the length of G_1_- and S-phase, suggesting that cell cycle progression may regulate their migratory behaviour.

### Notch signalling regulates TNC cell cycle progression

Our data show that Notch signalling allocates leader and follower identities, that cell cycle progression is necessary for TNC migration, and that leader and follower cells progress through the cell cycle at different rates. Does Notch signalling regulate cell cycle progression, thus differentiating leader from follower cells? To investigate this question, we measured the total length of the cell cycle and the length of each phase under control and Notch-inhibited conditions. While the total cell cycle span was not affected by Notch inhibition (Figure 7A), we found significant differences in the length of G_1_- and S-phase. Remarkably, leaders lose their characteristic cell cycle progression pattern and behave as followers, with a long G_1_ and a short S-phase (Figure 7B). Furthermore, Notch inhibition abolishes the size difference between migratory leader and follower cells, with all cells presenting the average follower’s area (Figure 7C-D). These data show that Notch activity defines TNC migratory identity by regulating cell cycle progression, cells with low Notch activity remain for longer in G_1_ behaving as followers. Interestingly, we noticed that Notch inhibition also changes the cell cycle behaviour of the followers’ population. While the followers’ average length of cell cycle phases is not altered, the dispersion of this population is significantly reduced, with standard deviations cut almost by half (from 2.7h to 1.42h for G_1_ and from 2.8h to 1.38h for S; Figure 7B). This prompted us to analyse the frequency distribution of cell cycle phases length. In control conditions, leader cells show a normal distribution with a single peak for G_1_- and S-phase, as expected for a homogeneous population. Followers, on the other hand, present a bimodal distribution, with the smaller peak coinciding with that of leader cells, and accounting for 26% of followers in G_1_- and 31% in S-phase (Figure 7E and F). Strikingly, these results fulfil the predictions of our *in-silico* model that best recapitulates TNC migration when chains are composed of leaders and followers in a 1:2 or 1:3 ratios. Furthermore, upon Notch inhibition the bimodal distribution of the follower population is lost, with all cells grouped at the major mean (Figure 7G and H). Taken together, our *in silico* and *in* vivo data demonstrate that the levels of Notch activity in TNC allocate migratory identity by controlling cell cycle progression, and indicate that migratory chains are formed of one leader cell every three followers.

**Figure 7.**
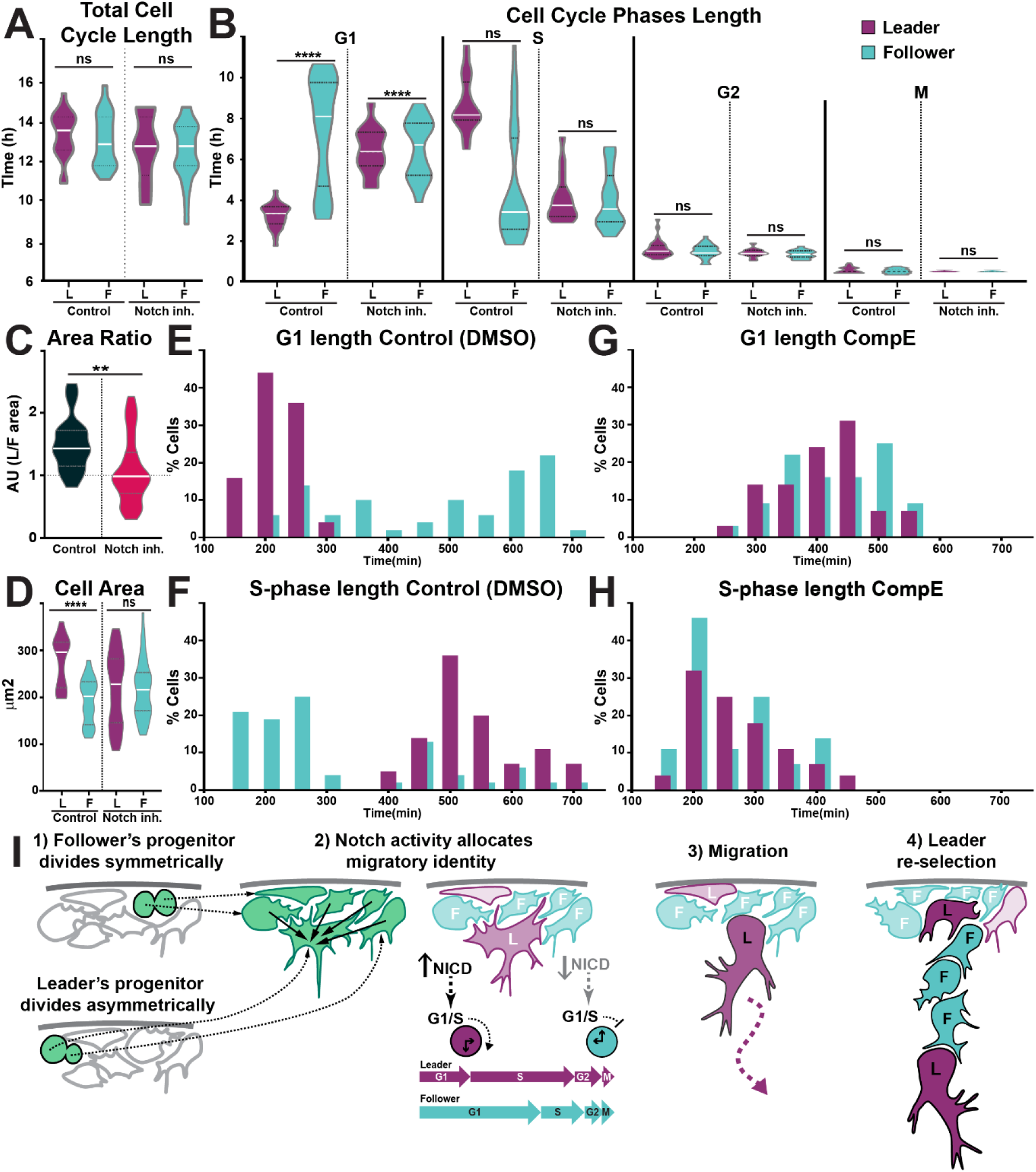
Notch signalling regulates TNC cell cycle progression. **A.** Quantification of the cell cycle total duration under control (DMSO, numbers as in Figure 6B) and Notch inhibition conditions (CompE, leaders n=17, followers n=22, 8 embryos; one-way ANOVA, p=0.1939). **B.** Quantification of the cell cycle phases duration under DMSO (numbers as in as in Figure 6C) and Notch inhibition conditions (CompE, leaders G_1_ n=29, S n=28, G2 n=25 and M n=25, 7 embryos; followers G_1_ n=32, S n=32, G2 n=30 and M n=30, 7 embryos; Brown-Forsythe and Welch ANOVA tests, all phases G_1_, S, G2 and M p>0.9999 between leaders and followers. **C.** Quantification of cell area under DMSO (leaders n=26, followers n=22, 6 embryos) and CompE conditions (leaders n=44, followers n=41, 7 embryos). Brown-Forsythe and Welch ANOVA tests, DMSO leaders vs followers p<0.0001, CompE All leaders vs followers p>0.9999. **D.** Quantification of cell area ratio (leaders/followers) under DMSO and Notch inhibited conditions (n as in E; Brown-Forsythe and Welch ANOVA tests, DMSO vs CompE All p=0.0157). **E-F**. Frequency distribution of G_1_- and S-phases durations in control conditions (DMSO; leaders: G_1_ n=45, S n=44, 11 embryos; followers: G_1_ n=50, S n=48, 11 embryos). **G-H**. Frequency distribution of G_1_- and S-phases durations in Notch inhibition conditions (CompE, leaders: G_1_ n=29, S n=28 7 embryos; followers: G_1_ n=32, S n=32, 7 embryos). **I**. Model of TNC acquisition through Notch signalling and cell cycle progression interactions.

## DISCUSSION

Collective migration plays an important role in embryogenesis, wound healing, and cancer. The acquisition of specific migratory identities has proven fundamental to angiogenesis, trachea development in Drosophila and cancer metastasis. TNC migrate collectively, forming chains with a leader cell at the front of the group that direct the migration, while follower cells form the body of the chain that trails the leaders. TNC leader and follower identities are established before migration initiation and remain fixed thereafter (Richardson et al., 2016). Herein, we have addressed the mechanism that establishes leader and follower identities and can propose the following model (Figure 7I): 1) premigratory TNC progenitors arise at the dorsal part of the neural tube. The leader’s progenitor divides asymmetrically giving rise to a large prospective leader cell and a small sibling that migrates as a follower. Other progenitors divide symmetrically giving rise to follower cells. 2) Interactions via Notch signalling results in the prospective leader cell accumulating higher levels of Notch activity. 3) The combination of high Notch activity and a larger cell size prompts the prospective leader cell to rapidly undergo the G_1_/S transition, entering S-phase and initiating migration earlier than its follower siblings, which are smaller and initiate migration whilst in G_1_. 4) Premigratory cells that have not been in contact with the prospective leader cell, or that have lost contact with it due to its ventral advance, maintain communication with surrounding premigratory TNC through Notch and undergo a new round of leader cell selection.

Notch signalling is a seemingly simple pathway that directly transduces receptor activation into changes in gene expression. Nevertheless, its outcomes in terms of cellular patterning are very diverse, from the generation of gene expression boundaries to temporal oscillations, or from the induction of similar fates in neighbouring cells to forcing adjacent cells into alternative fates. The latter function, known as lateral inhibition, is characterised by an intercellular negative feedback loop regulating the expression of Notch ligands. The activation of the Notch receptor in a “signal-receiving” cell leads to the downregulation of Notch ligands’ expression, making it less able to act as a “signal-sending” cell. The signature 2D patterning outcome of lateral inhibition is a mosaic of signal-sending cells with low Notch activity, surrounded by signal-receiving cells with high Notch levels. This is the case during the selection of sensory organ precursor cells in the epidermis of drosophila (Lewis, 1998), or the formation of the mosaic of hair cells and supporting cells in the sensory organs of the inner ear (Daudet and Żak, 2020). In general, however, lateral inhibition operates among cells subjected to extensive rearrangements and its patterning outcome is not a salt-and-pepper mosaic of cells (Bocci et al., 2020). For example, during angiogenesis, cells with low Notch signalling become tip or leaders, while cells with high Notch activity differentiate as stalk or followers (Phng and Gerhardt, 2009). In this context, leaders are interspaced by various numbers of followers. Several models have been proposed to explain how signal-sending (leader/tip) cells can exert a long-lasting or long-range inhibition on signal-receiving (follower/stalk) cells. These take into account the modulation of Notch signalling that arise from heterogeneity in Notch receptor levels, tension, Notch-regulators and interaction with other pathways (Bentley and Chakravartula, 2017; Koon et al., 2018; Kur et al., 2016; Venkatraman et al., 2016). Our data show that TNC deviate from the classical mosaic pattern, forming chains with one leader every two or three followers. Further study will be required to define whether the aforementioned mechanisms are responsible for this architecture.

In the case of the TNC, however, the most striking divergence from the classic lateral inhibition model (or indeed angiogenesis) is the fact that the leader cell identity is associated with higher intrinsic Notch activity. In other words, there are more signal-sending cells than signal-receiving cells. This apparent inversion in the ratio of the cell types produced is surprising. Explanation of this conundrum may arise from the fact that Notch lateral inhibition, dynamics and outcomes, can be modulated by “cis-inhibition”, a process whereby Notch ligands cell-autonomously interfere with the activation of Notch receptors (del Álamo et al., 2011; Bray, 2016). Computational models show that an increase in the strength of cis-inhibition can result in the inversion of the salt and pepper pattern (signal-sending to signal-receiving cells ratio), with the production of one cell with high Notch activity every three cells with low Notch levels (Formosa-Jordan and Ibañes, 2014), scenario that is congruent with the leader/follower ratio we observe in TNC. The detailed dynamics of lateral inhibition and whether cis-inhibition is at work in TNC remain to be investigated and will require direct visualisation at the single cell level of Notch activity in live embryos.

Our data show that active progression through the cell cycle is required for TNC migration. This is consistent with studies in chicken embryos, showing that NC continue cycling as they migrate, and that progression through G_1_/S is required for TNC delamination (Burstyn-Cohen and Kalcheim, 2002; Theveneau et al., 2007). Our data extend these findings by showing that leader and follower cells progress through the cell cycle at different rates. Leader cells, which are larger and more motile, initiate migration in S-phase and spend twice as long in this phase than followers. It is possible that these differences arise from the fact that leaders are larger than followers. It has been shown that the timing of G_1_/S transition depends on cell size and the dilution of the nuclear retinoblastoma protein (Zatulovskiy and Skotheim, 2020). Due to the larger volume of their cytoplasm leader cells could be primed for a rapid G_1_/S phase transition. The initiation of S-phase may in turn enhance leaders’ migratory characteristics through the interaction of cyclins and Cyclin/CDK inhibitors (CDKI) with small GTPases. Cyclin B and D, have been shown to phosphorylate cytoskeleton regulators, resulting in increased cell migration and tumour invasion (Blethrow et al., 2008; Chen et al., 2020; Chi et al., 2008; Hirota et al., 2000; Li et al., 2006; Manes et al., 2003; Song et al., 2008; Zhong et al., 2010). Furthermore, Rac1 activity, which is required for migration, oscillates during the cell cycle being highest at S-phase as cells are most invasive (Kagawa et al., 2013; Walmod et al., 2004). CDKIs, on the other hand, interact with RhoA and ROCK enhancing motility (Bendris et al., 2015; Creff and Besson, 2020; Yoon et al., 2012). Interestingly, enhanced motility increases actin branching, which in turn can accelerate the G_1_/S transition (Molinie et al., 2019). These factors could therefore generate a positive feedback loop in which slightly larger leader cells are prone to undergo the G_1_/S transition, in turn the activation of S-phase cyclins and CDKIs may enhance motility reinforcing S-phase initiation.

Our data also show that TNC cell cycle progression is under the control of Notch signalling. Upon Notch inhibition, all TNC present cell cycle phase lengths typical of follower cells. Notch has been shown to regulate cell cycle in a context-dependent manner. Depending on the cellular context, Notch has been shown to regulate the cell cycle through the transcriptional induction of Cyclin A and D, and the inhibition of CDKIs (Campa et al., 2008; Dabral et al., 2016; Ridgway et al., 2006; Rizzo et al., 2008; Rowan et al., 2008). Conversely, cell cycle progression can impact on Notch signalling. Notch activity is enhanced at the G_1_/S transition, while cells become refractory to Notch during G_2_/M (Ambros, 1999; Carrieri et al., 2019; Hunter et al., 2016; Nusser-Stein et al., 2012). Hence, the combination of large volumes and higher Notch activity levels could act synergistically to promote leaders’ G_1_/S transition.

In this study, we have uncovered new functional interactions between Notch signalling, cell cycle dynamics, and the migratory behaviour of leader and follower cells in the TNC. These complex and intricate interactions, which remain to be fully characterised at a molecular level, could apply to other cell types exhibiting collective migration. For example, studies in cancer cell lines have shown that activation or inhibition of Notch signalling hinders migration, similar to what we observe in TNC (Konen et al., 2017); while the maintenance of collective migration depends in on the regulation of cell proliferation during angiogenesis (Costa et al., 2016). In view of our work, it is important to revisit the assumption that migratory phenotypes are in conflict with cell cycle progression (Kohrman and Matus, 2017), and consider the possible implication for cancer therapies.

## ACKNOWLEDGMENTS

In memory of Julian Lewis. We are especially grateful to N. Daudet for his scientific and personal support. To the KCL fish facility staff, particularly to J. Glover. We are grateful to R. Kelsh, Y. Hinits and S. Wilson for sharing reagents. This project was funded by MRC G1000080/1, Royal Society 2010/R1 and Wellcome Trust 207630/Z/17/Z to CL; MR was supported by the Eunice Kennedy Shriver National Institute of Child Health & Human Development of the National Institutes of Health under awards T32HD055164 and F31HD097957. KB and DF-B were supported by the Francis Crick Institute core funding from CRUK (FC001751), MRC (FC001751), and Wellcome Trust (FC001751).

## AUTHOR CONTRIBUTIONS

ZA and CL designed the research. ZA, M A-G P, JR, TC and CL performed most of the experiments. ZA, DF-B, AG and CL analysed the experimental data. DF-B, CR and KB developed the computational model and performed the simulations. CL wrote the manuscript.

## STAR Methods

Key resources table

### Resource availability

Further information and requests for resources and reagents should be directed to and will be fulfilled by the lead contact, Claudia Linker

### Materials availability

Newly generated materials from this study are available by request from the Lead Contact, Claudia Linker

### Data and code availability

The code is being prepared for open sourcing on publication of this article.

### Experimental Models and Subject Details

#### Zebrafish lines and injections

Zebrafish were maintained in accordance with UK Home Office regulations UK Animals (Scientific Procedures) Act 1986, amended in 2013 under project license P70880F4C. Embryos were obtained from the following strains: *wild type, AB strain; Sox10:mG, Tg(−4.9sox10: Hsa.HIST1H2BJ-mCherry-2A-GLYPI-EGFP); Sox10:Fucci, Tg(−4.9sox10:mAGFP-gmnn-2A-mCherry-cdt1); hs:dnSu(H), vu21Tg (hsp70l:XdnSu(H)-myc); hs:Gal4, kca4Tg Tg(hsp70l:Gal4)1.5kca4 (1); UAS:NICD, kca3Tg Tg(UAS:myc-Notch1a-intra); Sox10:Kalt4, Tg(−4.9sox10: Hsa.HIST1H2BJ-mCherry-2A-Kalt4ER) and UAS:dnSu(H), Tg(UAS:dnSu(H)-myc); Tg(h2afva:GFP)kca13.* Embryos were selected based on anatomical/ developmental good health and the expression of fluorescent reporters when appropriate, split randomly between experimental groups and maintained at 28.5°C in E3 medium. Genotyping was performed by PCR of singe embryos after imaging when required (UAS:NICD; UAS:dnSu(H); hs:dnSu(H)). Injections were carried at 1-4 cell stage with 30pg of PCNA-GFP mRNA in a volume of 1nl. mRNA was synthesised from pCS2+ PCNA-GFP plasmid, kindly provided by C. Norden (IGC, Portugal), linearized with NotI and transcribed with the SP6 mMessage Machine Kit (Thermo Fisher Scientific, Cat#AM1340).

##### Live imaging and tracking

Imaging and analysis were carried as in Alhashem et al., 2021. In short, embryos were mounted in 1% agarose/E3 medium plus 40 μM Tricaine. Segments 6-12 were imaged in lateral views every 5 min from 16hpf for 16–18hr in an upright PerkinElmer Ultraview Vox system using a 40x water immersion objective. 70 μm z-stacks with 2 μm z-steps were obtained. Image stacks were corrected using Correct 3D Drift Fiji and single cell tracking performed with View5D Fiji plugin. Tracks were displayed using the MTrackJ and Manual Tracking Fiji plugins. Cell area measurements were done in Fiji using the freehand selection tool to draw around cell membranes in 3D stacks (as in Richardson et al., 2016). Cell speed measurements were calculated from 3D tracks using the following formula: ((SQRT((X1-X2)^2+(Y1-Y2)^2+(Z1-Z2)^2))/T)*60, where X, Y and Z are the physical coordinates and T is the time-step between time-lapse frames. Ventral distances were measured in a straight line from dorsal edge of the embryo to the cell position at the end of the movie. Cell directionality measurements were calculated using a previously published Excel macro (Gorelik and Gautreau, 2014). Total duration of the cell cycle was measured between two mitotic events. Cell cycle phases duration were measured using the characteristic nuclear pattern of PCNA-GFP, in movies where only TNC (expressing RFP and GFP) were shown using the following Fiji macro:

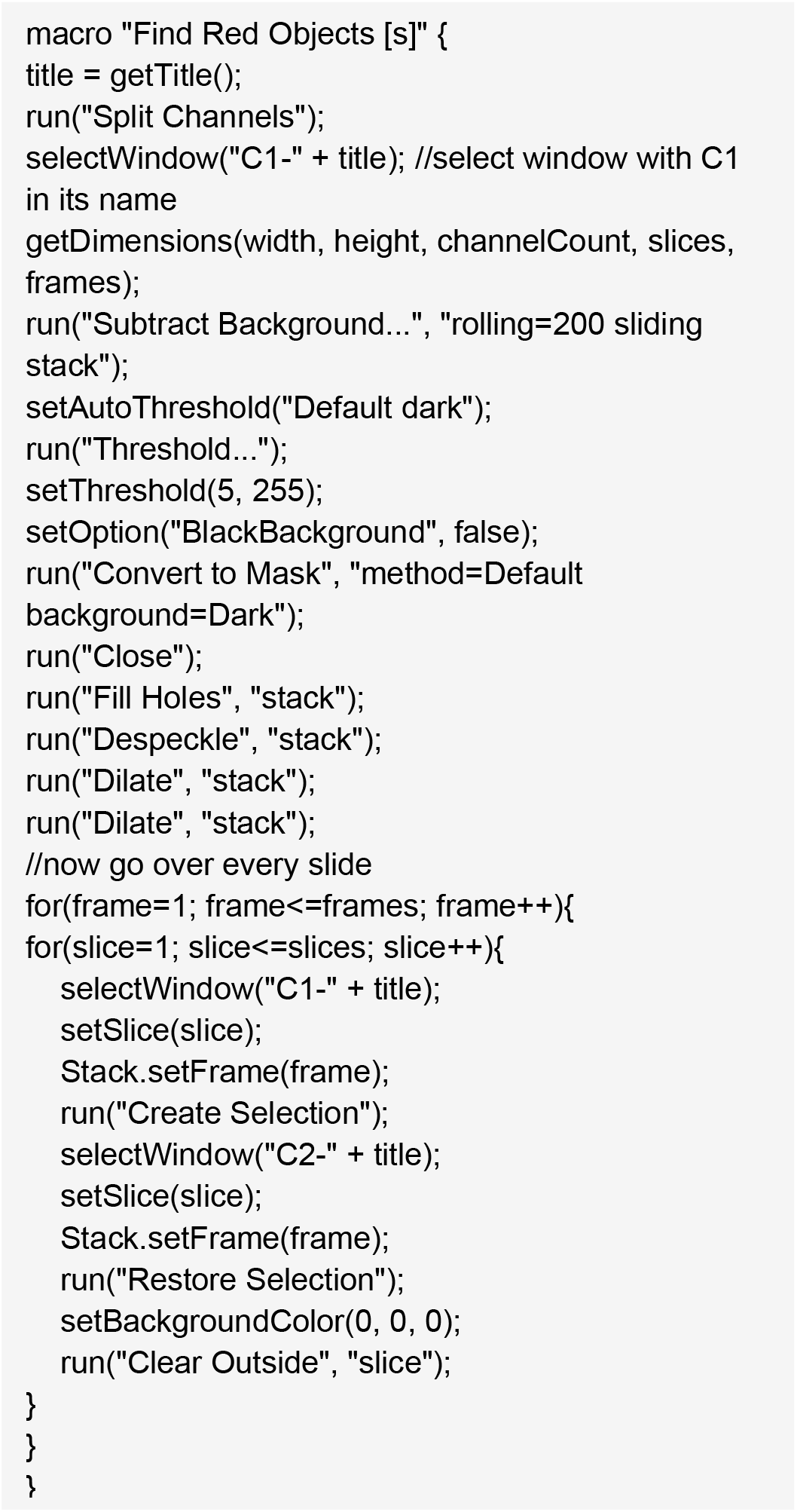

##### *Whole mount* in situ *hybridization, immunostaining, and sectioning*

The whole mount in situ hybridization protocol was adapted from https://wiki.zfin.org/display/prot/Whole-Mount+In+Situ+Hybridization. In short, embryos were fixed overnight (O/N) in 4% Paraformaldehyde (PFA), dehydrated in 100% methanol then rehydrated, digested with proteinase K for different times depending on the stage and pre-hybridised for 2h at 65°C. Riboprobes added, and embryos incubated at 65°C O/N. Probes were removed and embryos washed and equilibrated to PBS. Embryos were incubated in blocking solution for 2h and in anti-dig antibody O/N (Sigma-Aldrich Cat#41116161), washed 5×30min and NBT/BCIP colour reaction performed. Riboprobes for notch1a, deltaB, deltaD, her4, cb1045 were kindly provided by J. Lewis (CRUK); sox10 and foxd3 by R.N. Kelsh (University of Bath, UK); crestin, mbp, bdh, myoD, by S. Wilson (UCL, UK). After the in-situ colour development embryos were processed for sections, washed 5×10min with PBS, embedded in OCT, frozen by dipping the blocks in dry ice cold 70% ETOH, and sectioned to 12-15μm using a cryostat. Sections were defreezed at RT°C, incubated with blocking solution for 30min (10% goat serum, 2% BSA, 0.5% Triton, 10mM sodium azide in PBS) and in anti-GFP antibody ON 4°C (Millipore, Cat#06-896). Sections were washed with PBST 5×5min (0.5% Triton- PBS) and incubated with secondary antibody for 2h at RT°C, mounted in ProLong™ Gold Antifade Mountant (Molecular Probes Cat#P10144) and imaged. Wholemount antibody staining was performed in embryos fixed for 2h in 4% PFA, washed 4×10min, incubated in blocking solution for 2h and in primary antibodies O/N 4°C (anti-myc, Molecular Probes Cat#A21281; F59 and Znp1, Developmental Studies Hybridoma Bank; Acetylated tubulin, Sigma-Aldrich Cat#MABT868). Embryos were washed 5×30min, incubated in secondary antibodies O/N 4°C, washed 6×30min and mounted in 1% agarose for imaging. Imaging of sectioned and wholemount antibody-stained samples was performed in PerkinElmer Ultraview Vox system.

##### Drug treatments and gene expression induction

Embryos were treated by adding cell cycle inhibitors to the media from 11hpf and incubated for 3-12h at 28.5°C. 20μM Hydroxyurea (Sigma-Aldrich Cat#H8627), 300μM Aphidicolin (Sigma-Aldrich Cat#A0781), 100μM Genistein (Calbiochem Cat#345834), Teniposide (Sigma-Aldrich Cat#SML0609) or 1% DMSO as control (Sigma-Aldrich Cat#D8418). Notch signalling was inhibited at 11hpf by adding 100μM DAPT (Sigma-Aldrich, Cat#D5942-25MG) or 50μM of Compound E (Abcam Cat#ab142164), 1% DMSO was added as control conditions. Gene expression was induced by addition of 2.5μM of Tamoxifen (Sigma-Aldrich Cat#H7904) to the media at 11hpf of Sox10:Kalt4 embryos, or by heat shock at 11hpf in hs:Gal4 and hs:dnSu(H) embryos by changing the media to 39°C E3, followed by 1h incubation at this temperature, thereafter embryos were grown at 28.5°C to the desired stage.

##### Generation of UAS:dnSu(H) transgenic line

Using the Infusion cloning system (Takara) the following construct was inserted into the Ac/Ds vector (Chong-Morrison et al., 2018): 5xUAS sequence (Tol2Kit, http://tol2kit.genetics.utah.edu/index.php/Main_Page) flanked at the 3’ and 5’ ends by rabbit β-globin intron sequence. At the 3’ end GFP followed by SV40polyA sequence was cloned to generate the Ac/Ds dUAS:GFP vector. The cmlc2:egfp transgenesis marker (Tol2Kit) was cloned after GFP in the contralateral strand to prevent interaction between the UAS and the cmnl sequences. The *Xenopus* dnSu(H)-myc sequence (Latimer et al., 2005) was cloned into the Ac/Ds dUAS:GFP vector at the 5’ end of the 5xUAS sequence, followed by the SV40polyA sequence (Figure S4). Transgenesis was obtained by injecting Sox10:Kalt4 embryos with 1nl containing 50pg of DNA plus 30pg of Ac transposase mRNA at 1 cell stage. Embryos carrying the transgene were selected by heart GFP expression at 24hpf. Upon Gal4ER activation by tamoxifen dnSu(H)-myc protein was readily detected with anti-Myc antibody (Figure S4). GFP fluorescence driven by UAS was never observed.

##### Statistical analysis

All graphs and statistical analysis were carried out in GraphPad Prism 9. All numbers in the texts are average ± standard deviation. Every sample was tested for normality using the d’Agostino & Pearson, followed by the Shapiro-Wilk test. Samples that passed both tests were compared using either unpaired two-tailed *t*-test or one-way ANOVA. Those without a normal distribution were compared through a Mann-Whitney U test, Kruskal-Wallis test or Brown-Forsythe & Welch ANOVA tests. For all analyses, *p* values under 0.05 were deemed statistically significant, with \*\*\*\**p*<0.0001, \*\*\**p*<0.001, \*\**p*<0.01, and \**p*<0.05. Full statistical analysis of data in Figure 3 is presented in Table S1.

##### Computational model

The computational model used in this study is described in Methods S1.

### Supplementary Videos

**Video S1. Notch inhibition disrupts TNC migratory identity allocation. A-B**. Time lapse of Sox10:mG control (DMSO treated) embryo from 16-27hpf. **C-D**. Time lapse of Sox10:mG CompE treated embryo from 16-30hpf. Upper panels show fluorescent nuclei in grey and membranes in green. Lower panels show nuclei in grey, leaders tracked in magenta and followers in cyan. Arrowheads indicate leaders and arrows follower cells. Time in minutes. Related to Figures 2 and 3.

**Video S2. Notch gain and loss of function disrupts TNC migratory identity allocation.**

**A-B**. Time-lapse of control Sox10:Kalt4 embryo from 18-28.5hpf.

**C-D**. Time-lapse of Notch loss of function Sox10:Kalt4;UAS:dnSu(H) embryo from 18-27.9hpf.

**E-F**. Time-lapse of Notch gain of function, Sox10:Kalt4;UAS:NICD, embryo from 18-28.5hpf.

Upper panels show fluorescent nuclei in grey. Lower panels show nuclei in grey, leaders tracked in magenta and followers in cyan. Arrowheads indicate leaders, arrows follower cells. Time in minutes. Related to Figures 2 and 3.

**Video S3. *In silico* simulation of TNC chain migration.**

A. Simulation of a population with a single leader cell (Only 1L)

B. Simulation of a 1:1 leader follower ratio population (1L:1F)

C. Simulation of a 1:3 leader follower ratio population (1L:3F)

D. Simulation of a population composed only of follower cells (All followers)

E. Simulation of a population composed only of leader cells (All leaders) Leaders tracked in magenta and followers in cyan. Arrowheads indicate leaders, arrows follower cells. Time in minutes. Related to Figures 2 and 3.

**Video S4. Leader and follower cells present distinct division patterns.**

M>D time-lapse of Sox10:mG embryo from 16-23hpf, showing a leader cell dividing during migration.

D>M time-lapse of Sox10:mG from 16-28hpf, showing a follower cell dividing during before migration initiation.

Tracks before division in blue, after division in red and yellow. Arrows indicate divisions. Imaged from 16hpf to 28hpf. Related to Figure 4.

**Video S5. Cell cycle progression is required for TNC migration.**

Time-lapse of control (DMSO treated) Sox10:mG embryo from 16-23hpf, and Aphidicolin treated Sox10:mG embryo from 16hpf to 30hpf.

Leaders tracked in yellow, followers tracked in cyan and white. Time in minutes. Related to Figure 5.

**Video S6. Leader and follower cells initiate migration at different phases of the cell cycle.**

Representative time-lapse of Sox10:FUCCI from 16-18hpf showing leaders initiate migration in S-phase, while followers emigrate in G1.

Magenta arrowheads indicate the leader and its daughter cells; cyan arrowhead indicate follower cell. Time in minutes. Related to Figure 5.

**Video S7. PCNA-GFP reveals the cell cycle dynamics in TNC.**

Time-lapse of PCNA-GFP mRNA injected Sox10:Kalt4 embryo from 20-27.6hpf. Left raw image, right the same image showing only RFP^+^ TNC. Time in minutes. Related to Figures 6 and 7.

**Table S1. Statistical analysis of migratory parameters.**

Related to Figure 3.

### Supplementary Figures

**Figure S1:**
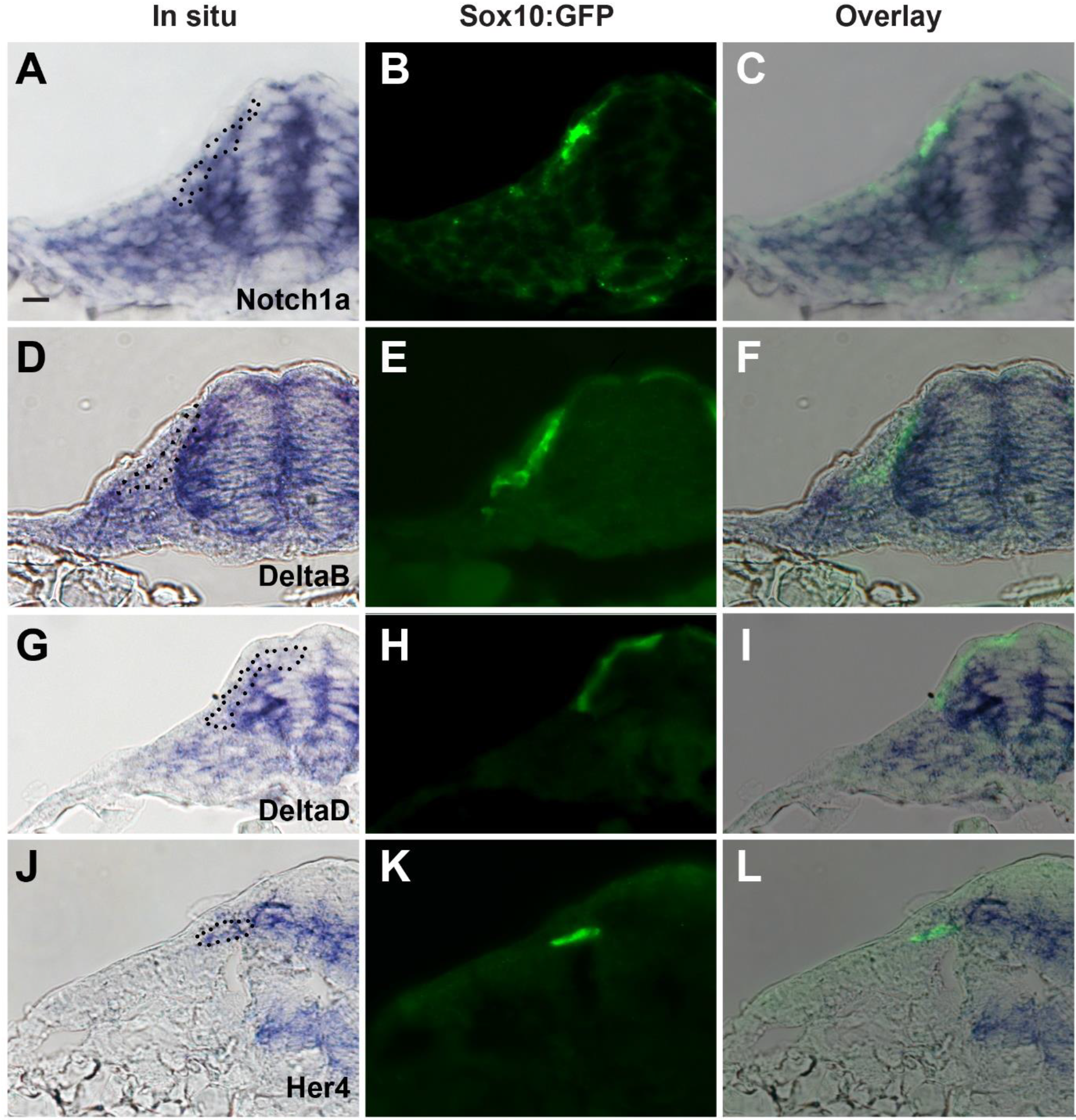
Expression of Notch signalling components during TNC migration. Transversal sections at trunk level of Sox10:GFP embryos showing the expression of: **A-C**. *Notch1a*, similar expression pattern was observed for *Notch1b*, *Notch2* and *Notch3* (not shown). **D-F**. *DeltaB* **G-I**. DeltaD, similar expression pattern was observed for *DeltaC* (not shown). **J-L**. *Her4* A, D, G and J bright field, B, E, H and K GFP-fluorescence and C, F, I and L overlay. Dotted black line in the brightfield frames indicates TNC cells seen in the fluorescent image. Related to Figure 1.

**Figure S2:**
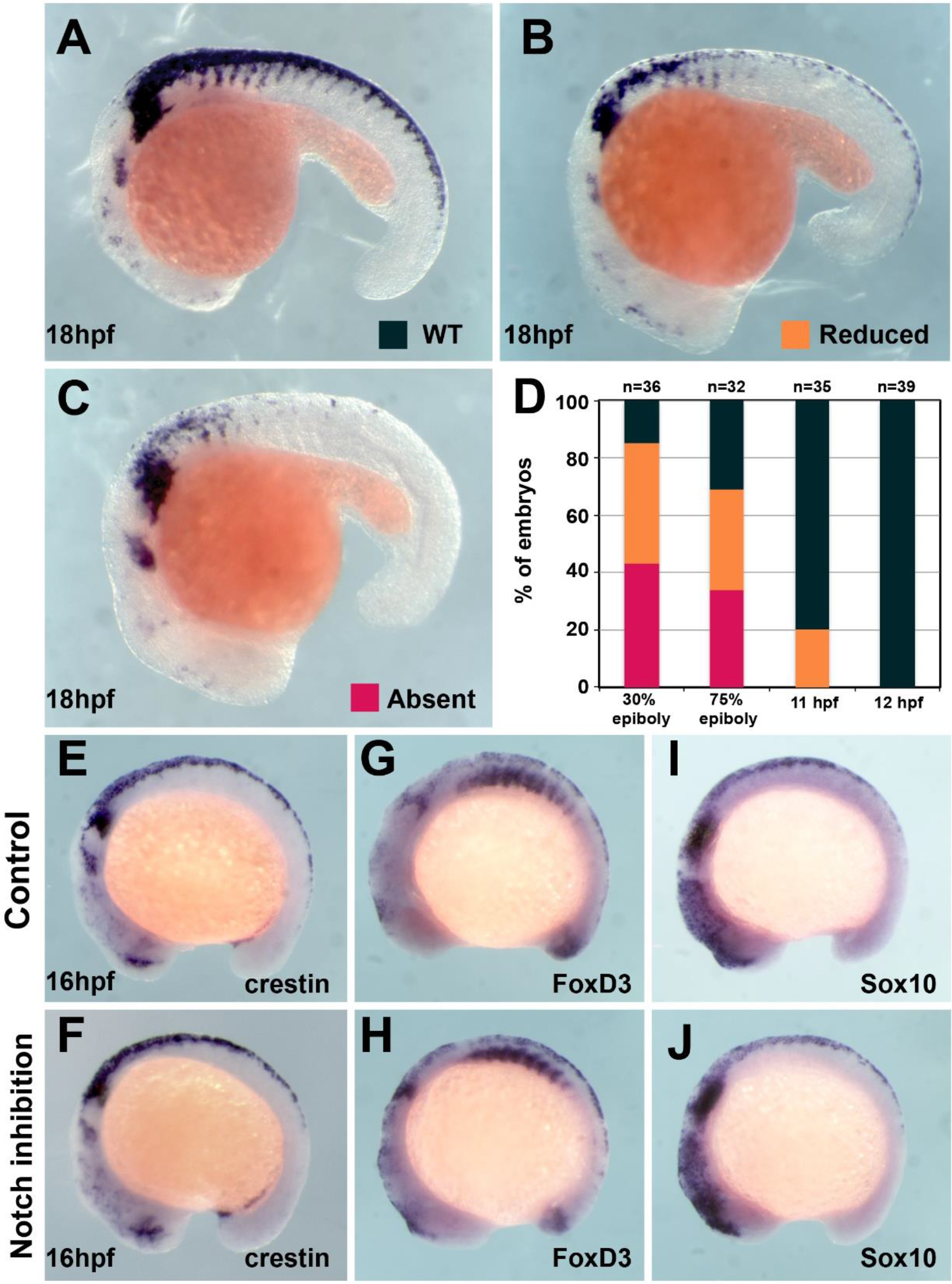
TNC induction is independent of Notch signalling after 12hpf. **A**. *Crestin* in situ hybridisation in wildtype (WT) embryo at 18hpf. **B-C**. *Crestin* in situ hybridisation in DAPT treated embryos: (B) reduced or (C) absent TNC. **D**. Quantification of the *Crestin* expression phenotypes upon DAPT treatment (phenotypes: WT black; reduced orange; absent red; 30% epiboly n=38, 75% n=32, 11hpf n=35, 12hpf n=39). **E-J**. In situ hybridisation for NC markers in control (DMSO) and DAPT treated embryos from 12-16hpf. (E and F) *Crestin* (DMSO n=32, DAPT n=38), (G and H) *FoxD3* (DMSO n=16, DAPT n=35) and (I and J) *Sox10* (DMSO n=27, DAPT n=29). Anterior to the left, dorsal top. Related to Figure 1.

**Figure S3:**
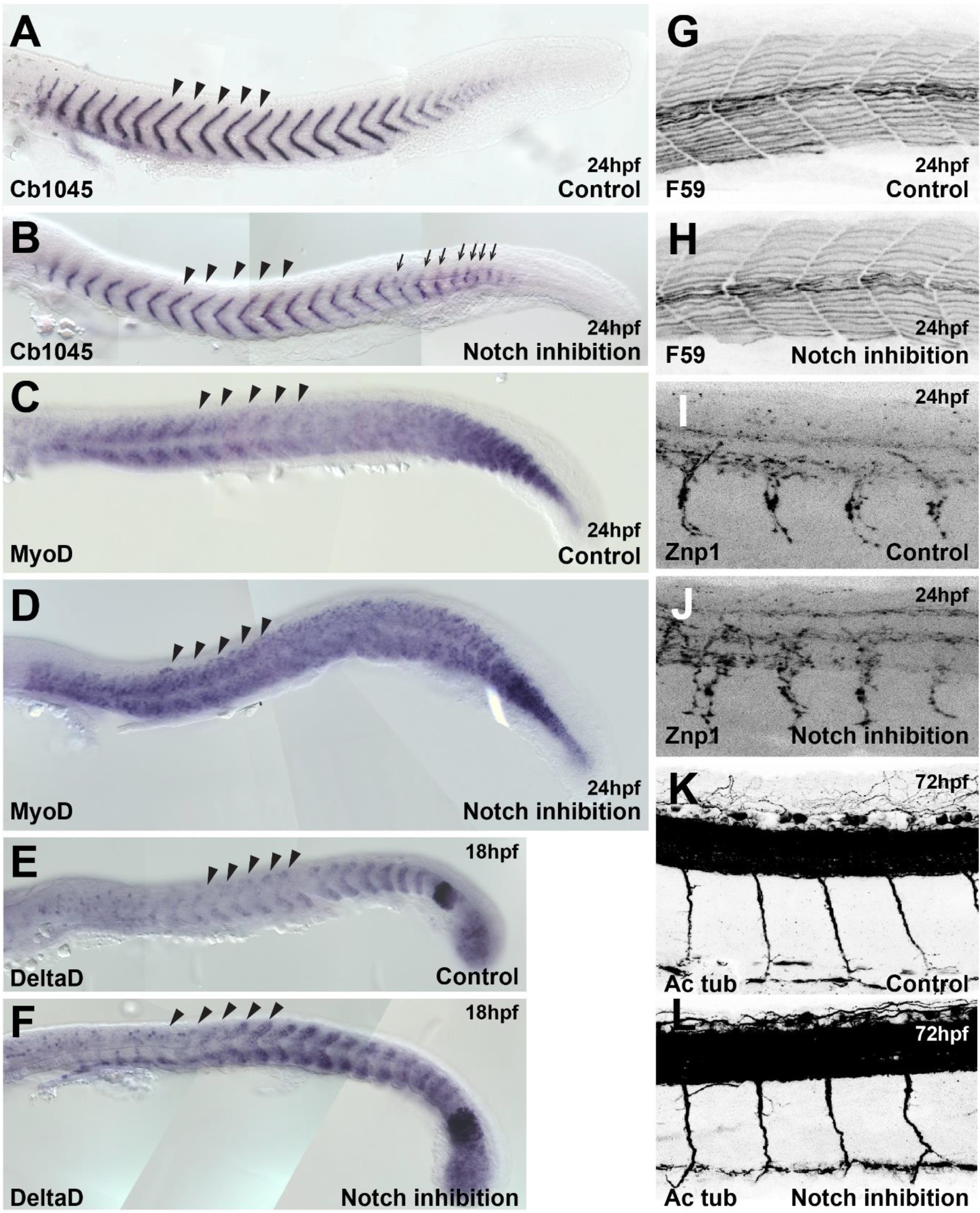
Somites and neural tissue formation are not altered by Notch inhibition. **A-B**. *cb1045* expression showing somite boundaries upon (A) control (DMSO, n=23) and (B) DAPT (n=30) treatment. Arrows indicate segmentation defects. **C-D**. *MyoD* expression showing muscle progenitors upon (C) control (DMSO, n=47) and (D) DAPT (n=45) treatment. **E-F**. *DeltaD* expression showing somite polarity upon (E) control (DMSO, n=25) and (F) DAPT (n=30) treatment. **G-H**. Antibody staining for heavy myosin (F59), showing differentiating muscle cells, upon (G) control (DMSO n=37) and (H) DAPT (n=32) treatment. **I-J**. Antibody staining for Znp1, showing neurons, upon (I) control (DMSO n=35) and (J) DAPT (n=42) treatment. **K-L**. Antibody staining for acetylated tubulin (Ac. Tub), showing neurons, upon (K) control (DMSO n=20) and (L) DAPT (n=27) treatment. Arrowheads indicate the level at which TNC migration was analysed Anterior to the left, dorsal top. Related to Figure 2 and 3.

**Figure S4:**
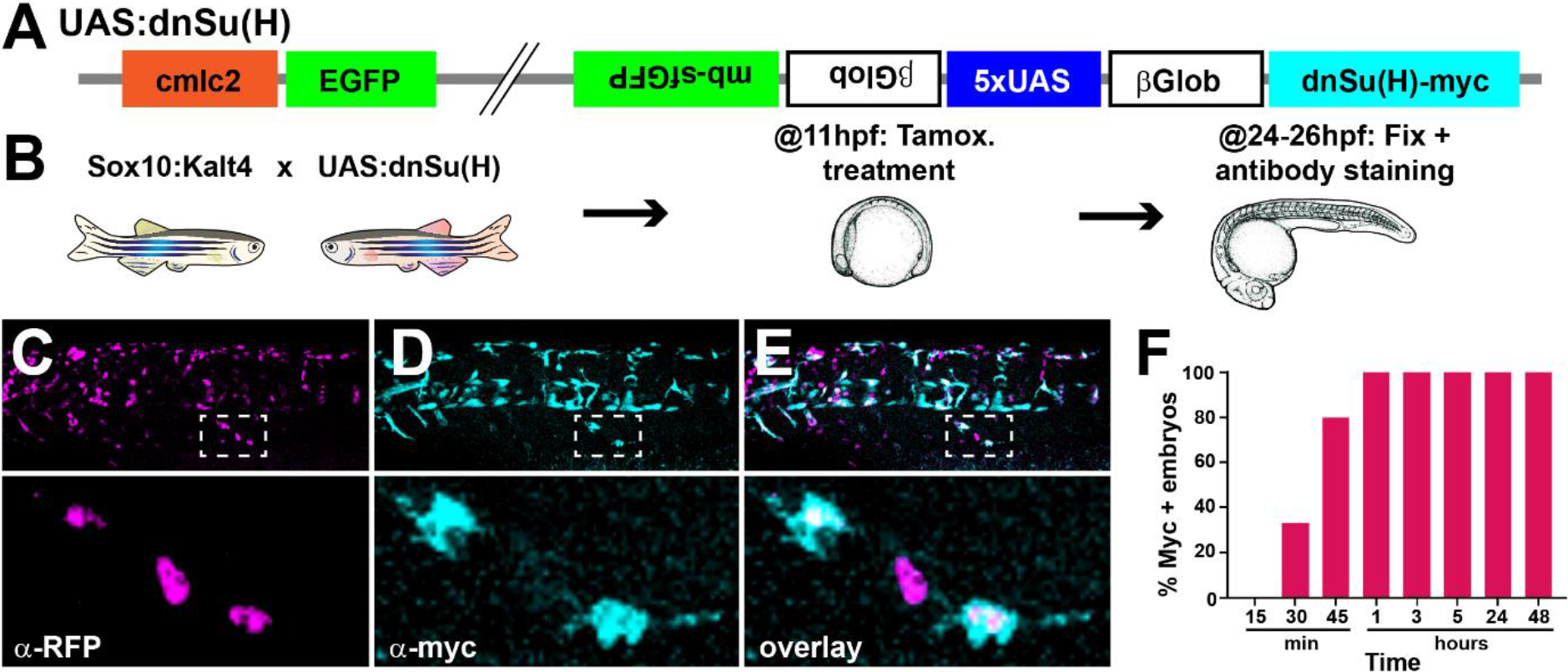
UAS:dnSu(H) transgenic line. **A.** Diagram of the construct used to generate the UAS:dnSu(H) line. **B.** Scheme of protocol used. **C-E.** Trunk region of a Sox10:Kalt4;UAS:dnSu(H) embryo treated with tamoxifen from 11-24hpf and immunostained for (C) RFP and (D) myc. (E) overlay. Dotted squares indicate enlargement. **F.** Number of embryos expressing the UAS driven (myc+) after tamoxifen treatment from 11hpf for different times (15’ n=20, 30’ n=27, 45’ n=25, 1h n=22, 3h n=18, 5h n=20, 24h n=14, 48h n=10). Related to Figure 2 and 3.

**Figure S5:**
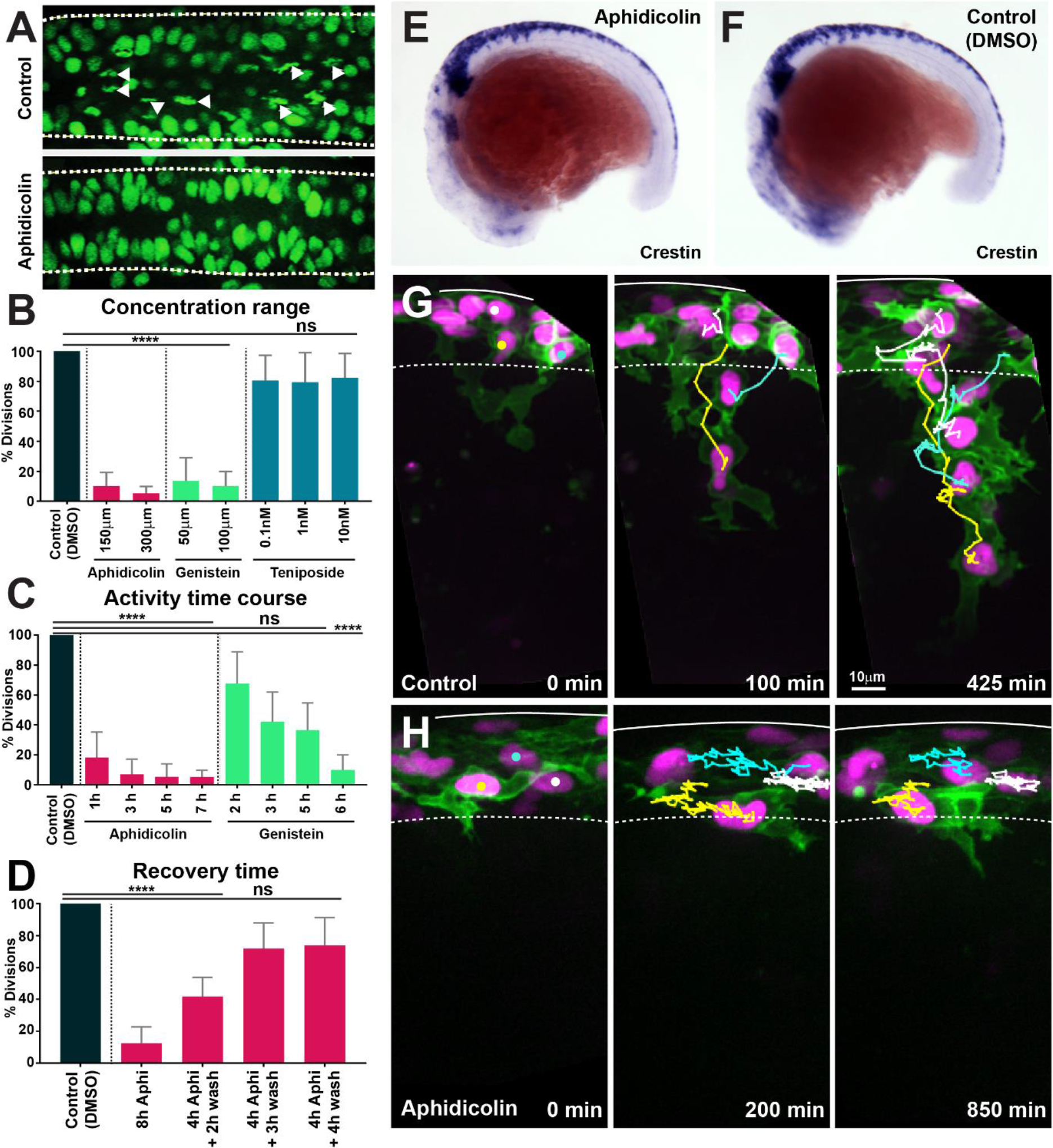
Cell cycle inhibitors drugs working conditions. **A.** Confocal images showing nuclei and mitotic figures in Control (DMSO) and aphidicolin treated H2A:H2A-GFP embryos. Arrowheads indicate mitotic figures; dashed lines mark the neural tube borders. Dorsal view, anterior to the left. **B.** Percentage of mitotic figures respect to Control (DMSO) in embryos treated with different concentrations of cell cycle inhibitors (Kruskal-Wallis test, p<0.0001, aphidicolin n=20 and genistein n=32; teniposide p>0.9999 n=27). **C.** Time-course of cell cycle drugs effect (Aphi: Aphidicolin; Kruskal-Wallis test, Control vs 1h Aphi p=0.0007; control vs 3h, 5h and 7h Aphi p<0.0001; control vs 2h, 3h and 5h Geni p>0.0892; control vs 6h Geni p<0.0001; control n=62 embryos; Aphi 1h n=16, Aphi 3h n=15, Aphi 5h n=16, Aphi 7h n=15; Geni 2h n=15, Geni 3h n=16, Geni 5h n=17, Geni 6h n=15). **D.** Quantification of cell cycle recovery times following aphidicolin removal (Aphi: Aphidicolin; control n=21; Aphi 8 hours n=18, Aphi 4+2h wash n=15, 4+3h wash n=15 and 4+4h wash n=15 embryos; One-way ANOVA, Control vs 8h and 4+2h wash *p*<0.0001; control vs 4+3h wash and 4+4h wash *p*>0.0851). **E-F**. *Crestin* expression in 16hpf embryos upon (E) aphidicolin and (F) DMSO treatment from 11hpf. Anterior to the left, dorsal top. **G-H**. Selected frames of *in vivo* imaging from Sox10:mG embryos showing cell tracks under (G) control (DMSO) 16-28hpf and (H) aphidicolin 16-33hpf treated. Solid line indicates the dorsal midline, dashed line the premigratory area; time in minutes. Related to Figure 5.

**Figure S6:**
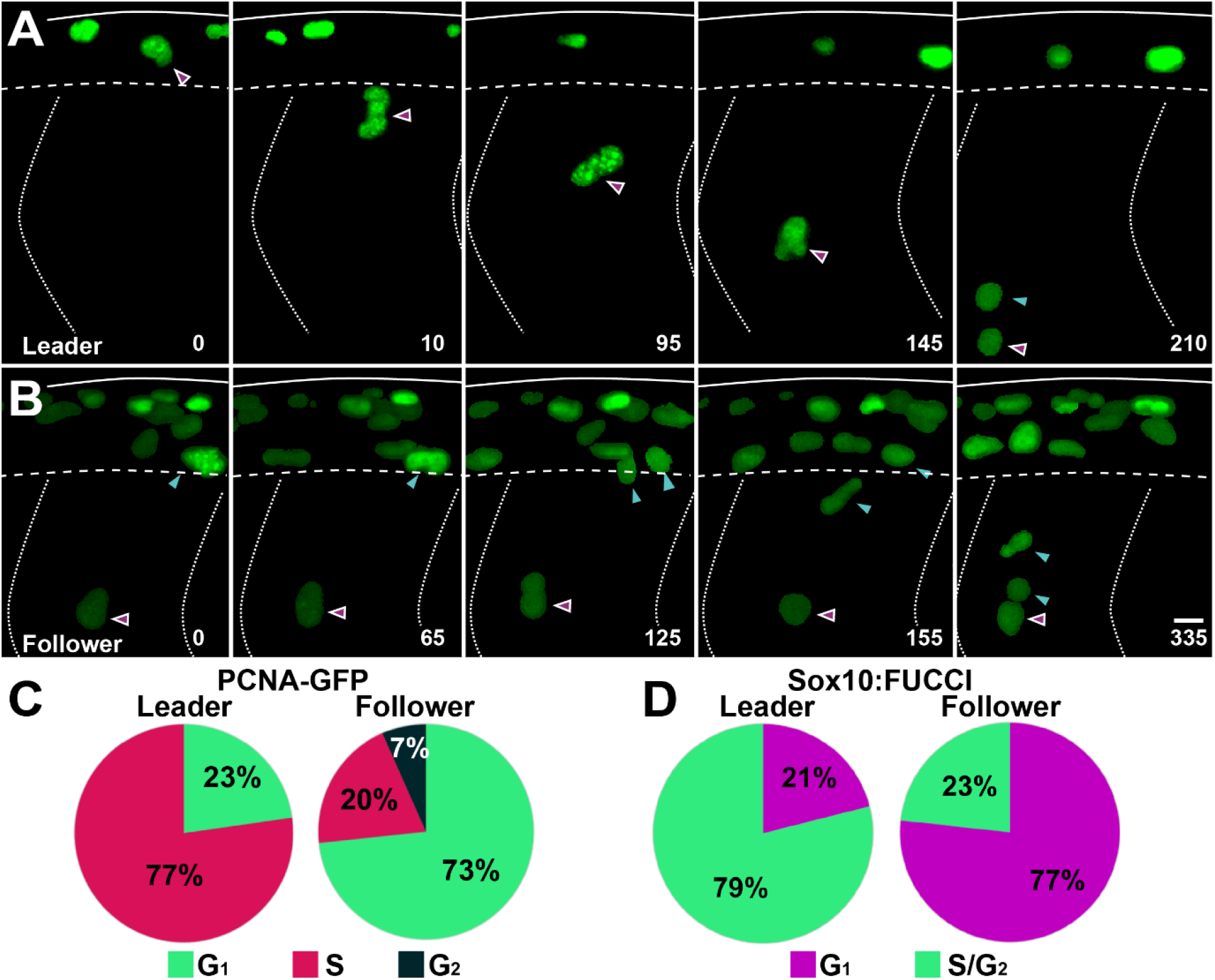
Leader and follower cells initiate migration at distinct cell cycle phases. **A-B**. Selected frames of *in vivo* imaging from Sox10:Kalt4 embryos injected with PCNA-GFP mRNA, showing PCNA localization in RFP labelled TNC. (A) Leader cell initiates migration in S-phase. (B) Follower cell divides and initiate migration in G_1_. Solid lines indicate embryo dorsal border, dotted lines the somites borders, segmented line the premigratory ventral border. Time in minutes. Anterior to the left, dorsal up. **C.** Quantification of the cell cycle phase at which cells initiate migration in PCNA-GFP mRNA injected embryos (leaders n=22, 10 embryos; followers n=45, 10 embryos). **D.** Quantification of the cell cycle phase at which cells initiate migration in Sox10:FUCCI embryos (leaders n=38, 4 embryos; followers n=43, 4 embryos). Related to Figure 6.

