## Supplementary material for "Notch controls the cell cycle to define leader versus follower identities during collective cell migration": Methods S1

### Methods S1: Computer Modelling Methods

A minimal discrete element model of TNC migration was developed in which each cell is modelled as an infinitesimal particle moving in 2D space. A network of neighbours within the particle system is identified by a Delaunay triangulation (Figure 1a).

Equation 1: $V(r) = D_e\left(e^{-2a(r-r_e)} - 2e^{-a(r-r_e)}\right),\ \ a = \sqrt{k/\ 2D_e}$

Thus defined, the system exhibits Brownian dynamics as described by the over-damped Langevin equation [Equation 2], such that the velocity of each particle is proportional to the resultant force applied to it ($\nabla V$), plus a stochastic component ($\zeta$).

Equation 2: $\underline{\dot{x}} = -\frac{\Delta V}{\gamma} + \underline{\zeta}$

Components of the resultant force on each cell arise from cell-cell interactions, cell-boundary interactions, and cell autonomous motio

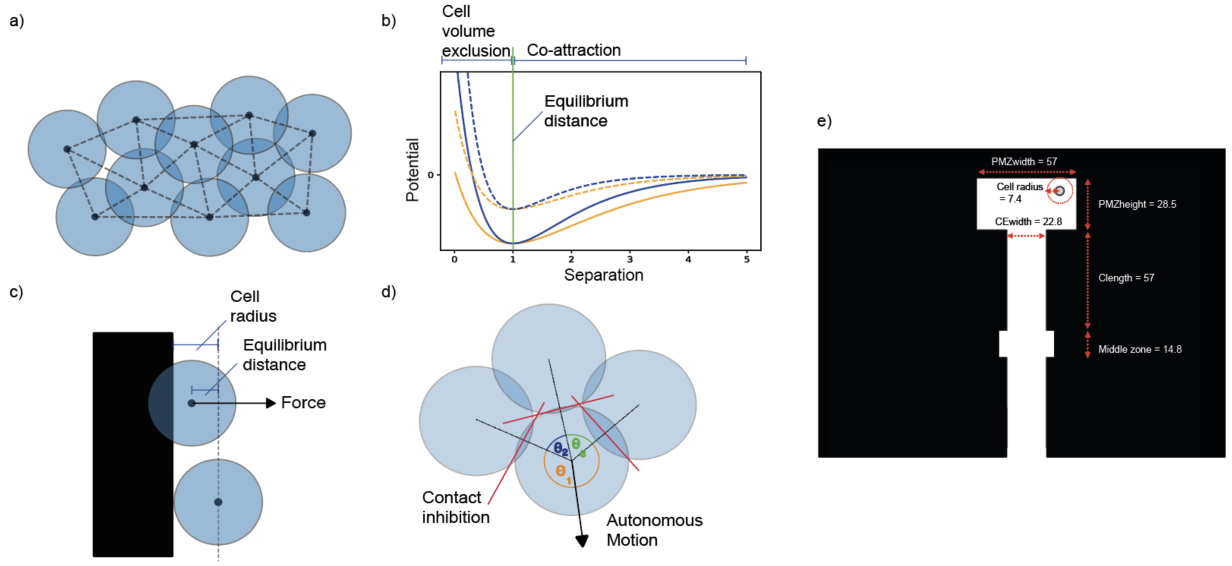


**Figure 1. Description of model mechanisms and configuration**

a) Diagram of 10 cells modelled as infinitesimal particles, with Delaunay triangulation showing nearest neighbours and circles showing typical cell radii around each particle.

b) Morse potential for low volume exclusion k (orange), high cell volume exclusion (blue), high energy depth De (solid line), and low energy depth (dashed line). The portion of the curve that relates to repulsion is distinguished from the portion that relates to attraction by the vertical green line.

c) Demonstrating calculation of force component from a boundary. When the centre point of a cell moves within a cell radius of the boundary, the cell experiences a force perpendicular to and away from the boundary with magnitude determined by a Morse potential and with offset from equilibrium distance.

d) Demonstrating calculation of cell polarisation. Adjacent nearest neighbours of a cell subtend angles θ1, θ2, and θ3 around the cell centre. The direction of polarisation, and hence autonomous motion, bisects θ3, the largest such angle. Forces on each cell arise from interactions between neighbouring particles. These interactions are defined by a Morse potential [Morse, 1929], a function of the separation between particles, and parameterised by an equilibrium separation (re), approximate spring constant (k), and energy depth (De) [Equation 1, Figure 1b]. These parameters model the typical radius of a cell, its volume exclusion, and chemoattractive magnitude (“co-attraction”).

e) Dimensions of the model. White space represents empty space where cells can move freely, black space is space where cells cannot move due to boundaries. Horizontal movement is restricted while moving down the chain except for in the middle zone (for values associated with these parameters see table 1).

**a. Tissue environment (boundary)**

Cells move into permissive space between the neural tube/notochord and the somites. Boundary locations are specified before any simulation. The boundary is implemented as a region of space that applies strong repulsion to nearby cells (Figure 1e). Any cell that moves within a cell radius of the boundary experiences a force given by the gradient of the same Morse potential used in cell-cell interactions, such that the repulsion of any cell from the boundary depends upon the cell volume exclusion and increases exponentially as the cell approaches the boundary (Figure 1e).

The size and shape of the boundary represent a space for the pre-migratory cells at the top, a space in the middle where the notochord and neural tube meet (midline) and a vertical space where the chain can proceed downwards.  The dimensions of the environment boundary were calibrated to *in vivo* measurements (Figure 1e – showing micron scale dimensions on the boundary).

The system is setup in a 'T' shape, which is interrupted in the middle by a space of horizontal mobility, because *in vivo* cells regularly move into this space. The wider region at the top represents the premigratory zone (PMZ) at the top of each migratory chain. Cells are able to filter in from the sides to mimic the continuous clustering of cells above migration chains.

**b. Cell properties/ behaviours**

b.1. Contact inhibition and autonomous motion

Cells exhibit autonomous motion in a direction determined by their internal polarisation. This polarisation is influenced by interaction with the cell's neighbours, such that the cell will try to move into empty space. We introduce contact inhibition into the model as a term in the Langevin equation [Equation 2], with magnitude determined by a user-defined parameter. The direction of autonomous magnitude for a given cell is found by identifying all adjacent nearest neighbours surrounding the cell, calculating the angle subtended by each adjacent pair, and bisecting the largest such angle (Figure 1d). The magnitude of this autonomous velocity component is proportional to the user-defined parameter (aMag) and the square of the maximum subtended angle, representing the combined effect of greater polarisation and more free space to move into. Any cell that moves beyond a threshold distance from its nearest neighbour will stop autonomous motion, modelling the loss of polarisation when losing contact with neighbouring cells.

b.2. Cell volume exclusion

Cells exhibit volume exclusion (two cells repel from one another if they get closer than an equilibrium distance). This simply models how two cells cannot occupy the same space at the same time. The extent to which volume exclusion is exhibited can be thought of as the level of cell stiffness. Low $k$ means cells are squishier. This is modelled using the $k$ term in the Morse potential calculation [Equation 1].

b.3. Co-attraction (co-A)

When cells drift more than the equilibrium distance apart, they are drawn back toward their neighbours with a force calculated by the Morse potential curve [Equation 1].

**c. Migratory Identity**

Leader and follower migratory identities were allocated to cells according to the order in which these enter the chain. That is, the first cell becomes leader then the next X many cells become follower cells before the cell after that becomes leader. A sensitivity analysis on leader cells frequency was performed by spacing parameter S.

**d. Simulation procedure**

The simulation follows the process steps in Figure 2 and was simulated on CAMP – the Francis Crick Institute's Linux-based high-performance computing system. Parameter combination/ experimental condition pairs were run 100 times in parallel across 10 nodes.

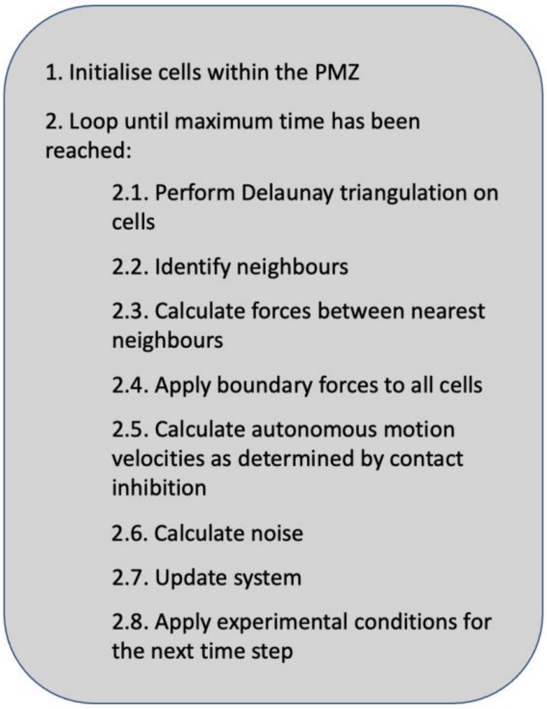


**Figure 2. Model pseudocode overview.**
A predefined number of cells is initialised in the PMZ. Thereafter, the system enters a loop for every time step up to tmax. In this loop, forces are linearly summed to obtain each cell's velocity vector for that time step:
1) a Delaunay triangulation is performed on cells.
2) Each cell's nearest neighbours are identified.
3) Local forces between cells are calculated according to the Morse potential (Figure 1b).
4) Boundary forces are applied to each cell.
5) Autonomous motion and contact inhibition are calculated for each cell.
6) Gaussian noise is added to each cell's velocity vector.
7) The system's clock is updated, as well as each cell's position.
8) Experimental conditions are applied for the next time step (e.g., giving certain cells leader qualities).

**e. Parameterisation**

Time was calibrated as follows: *in vivo* control cells tend to migrate to approximately 120 microns from dorsal midline on average (Figure 3F main text). The total time of migration is on average 11.64h long (~700 minutes). In 2000 timesteps, the control case (with differential CIL, heterogenous migratory identities and S = “1:3”) also migrates to approximately 120mm. We gathered data every 20 timesteps, which means in our simulation movies there are 100 frames (i.e., 1 frame = 7 minutes).

Where possible parameters were calibrated to values measured *in vivo* (Table 1). Model specific parameters unable to be linked directly to *in vivo* values were set to values that produced realistic bounds of behaviour.

**Table 1. Simulation parameters, description, range and source**

| Name | Description | Range | Optimised setting | Units | Source |
|---|---|---|---|---|---|
| PMZ width | Horizontal space of the premigratory zone | 57.0 | 57.0 | mm | Measurem nt |
| PMZ height | Vertical space in the premigratory zone | 28.5 | 28.5 | mm | Measurem nt |
| CE width | Horizontal width in the migratory chain | 22.8 | 22.8 | mm | Model specific |
| MZ ratio | Vertical space around the midpoint relative to the heigh of the PMZ | 0.5 | 0.5 | Units | Model specific |
| Cell radius | Interaction radius of cell radiu was inferred assuming cells were perfect spheres, based c volumetric measurements (Richardson et al., 2016) | 7.4 | 7.4 | mm | Measurem nt |
| Nc | Number of cells | 18 | 18 | Number | Measurem nt |
| $\zeta$ | Magnitude of stochastic component. Term of the Langevin equation, which controls random cell moveme magnitude | 0.035 | 0.035 | Units | Model specific |
| $\gamma$ | Overdamped Langevin equation drag factor. | 1 | 1 | Units | Model specific |
| S | Leader spacing- number of follower cells between leader cells in migration | {0, 1, 2, 3, ∞} | 3 | Number | Calibrated |
| Follower k | Spring constant near equilibrium (parameter of Morse potential) for follower type cells. This can be though of as the cell volume exclusio of the cells. High k means tha cells are stiffer. | Low: [0.01] Medium: [0.02] High: [0.03] | 0.01 | Units | Calibrated |
| Leader k | As above but for leader type cells. | Low: [0.01] Medium: [0.02] High: [0.03] | 0.02 | Units | Calibrated |
| Follower De | Depth of potential well (parameter of Morse potential Greater De means greater | Low: [3e-05] | 3e-05 | Units | Calibrated |

| | | | | | |
|---|---|---|---|---|---|
| | range of co-attraction. This ca be thought of as the amount c chemotactic attraction signal released by each cell. | Medium: [6e 05]<br><br>High: [9e-05] | | | |
| Leader De | As above but for leader type cells. | Low: [3e-05]<br><br>Medium: [6e 05]<br><br>High: [9e-05] | 6e-05 | Units | Calibrated |
| Follower aMag | Magnitude of autonomous cell velocity. In the model's implementation of contact inhibition, cells move into the widest open space. This parameter modulates the velocity with which they move into this space. | Low: [1.1e-07]<br><br>Medium: [1.56e-06]<br><br>High: [3e-06] | 1.1e-07 | Units | Calibrated |
| Leader aMa | As above but for leader type cells. | Low: [1.1e-07]<br><br>Medium: [1.56e-06]<br><br>High: [3e-06] | 3e-06 | Units | Calibrated |
| Interaction threshold | Multiples of cell radii beyond which neighbours no longer cause polarisation by contact inhibition. | 1 | 1 | Units | Model specific |
| T max | Total run time in arbitrary units. | 2000 | 2000 | Units | Model specific |
| dt | Time interval between iterations. | 0.1 | 0.1 | Units | Model specific |
| Output interval | Time interval between data outputs. | 10 | 10 | Units | Model specific |

**f. Sensitivity analysis**

In the grid search calibration approach we fixed follower cells properties to be at their low levels. Next, we looked at how changes to leader cell physical properties affected ventral distance (Figure 3). This shows a strong effect in leader aMag, whereby low leader aMag resulted in cells not migrating much beyond 100 microns no matter the level of co-attraction or cell volume exclusion. aMag had to be varied across a wider range to see a clear effect. Through this, aMag has a dominating effect on ventral distance: higher aMag is associated with higher ventral distance.

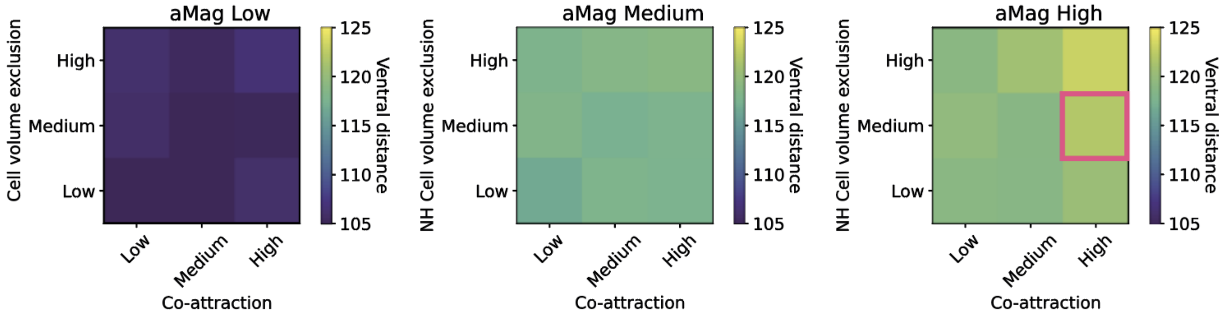


**Figure 3. Calibration on final position of the furthest travelling cell in microns.**
The optimal distance is 120 microns which is shown by the pink square. Ventral distance increases with increases in cell volume exclusion and co-attraction, which is most apparent in the rightmost heatmap.
